# In heart failure reactivation of RNA-binding proteins drives the transcriptome into a fetal state

**DOI:** 10.1101/2021.04.30.442191

**Authors:** Matteo D’Antonio, Jennifer P. Nguyen, Timothy D. Arthur, Hiroko Matsui, Margaret K.R. Donovan, Agnieszka D’Antonio-Chronowska, Kelly A. Frazer

**Author notes:** The authors wish it to be known that, in their opinion, the first two authors should be regarded as joint First Authors.

## Abstract

Transcriptome-wide expression changes occur during heart failure, including reactivation of fetal-specific isoforms. However, the underlying molecular mechanisms and the extent to which a fetal gene program switch occurs remains unclear. Limitations hindering transcriptome-wide analyses of alternative splicing differences (i.e. isoform switching) in cardiovascular system (CVS) tissues between fetal and adult (healthy and diseased) stages have included both cellular heterogeneity across bulk RNA-seq samples and limited availability of fetal tissue for research. To overcome these limitations, we have deconvoluted the cellular compositions of 996 RNA-seq samples representing heart failure, healthy adult (heart and arteria), and fetal-like (iPSC-derived cardiovascular progenitor cells) CVS tissues. Comparison of the expression profiles revealed that RNA-binding proteins (RBPs) are highly overexpressed in fetal-like compared with healthy adult and are reactivated in heart failure, which results in expression of thousands fetal-specific isoforms. Of note, isoforms for 20 different RBPs were among those that reverted in heart failure to the fetal-like expression pattern. We determined that, compared with adult-specific isoforms, fetal-specific isoforms are more likely to bind RBPs, have canonical sequences at their splice sites and encode proteins with more functions. Our findings suggest targeting RBP fetal-specific isoforms could result in novel therapeutics for heart failure.

## Introduction

RNA binding proteins (RBPs) play a pivotal role in regulating alternative splicing outcomes (i.e. relative abundance of different alternatively splice isoforms) and hence are major drivers of transcriptomic diversity during cardiac development and between adult heathy and diseased states ^1,2^. Several genes with pivotal roles in cardiac development and function, including *SCN5A*, *TNNT2*, *ABLIM1* and *TTN*, have been shown to express different isoforms in the fetal and adult stages; likewise, multiple RBPs (including *CELF1*, *HNRNPL*, *RBFOX2*, *RBM24*, *MBNL1* and *RBFOX1* ^3–9^) have been found to have stage-specific expression. In heart failure, the adult heart reverts to a fetal-like metabolic state and oxygen consumption ^10,11^ and has been shown to reactivate expression of several fetal-specific genes, including *ACTA1* ^12^ and fetal type cardiac ion channels ^13^, and suppress several adult-specific genes such as *SERCA2* ^14^ and *MYH6* ^15^. While previous studies suggest that alternative splicing plays an important role during heart development ^16,17^ and a return to the “fetal gene program” occurs during heart failure, transcriptome-wide alternative splicing changes during cardiac development and disease, and how RBPs regulate these changes, have yet to be investigated.

Until now, several limitations have hindered transcriptome-wide analyses of alternative splicing differences (i.e. isoform switching) in cardiovascular system (CVS) tissues between fetal and adult (healthy and diseased) stages. CVS tissues (heart and arteria) are composed of multiple cells types, and different samples of the same tissue type are comprised of different proportions of these cell types ^18,19^. Heterogeneity across bulk RNA-seq samples of the same tissue type limits the power to identify gene and isoform expression differences between different tissues and development stages. Furthermore, bulk RNA-seq experiments average gene expression across the population of cells in a given sample and thereby prevent the ability to capture isoform expression variability across different cell types. While in theory single cell transcriptomics may be more indicative than bulk RNA-seq for this type of analysis, most of the recently developed technologies fail to assess the full transcriptome, as they rely on the expression of only the 3’ end of each transcript and, therefore, cannot be used to investigate isoform expression. Finally, the limited availability of human fetal heart tissues for research makes it impossible to obtain the large sample sizes required for the statistical power to perform transcriptome-wide analyses of the re-expression of fetal-specific isoforms.

To overcome these limitations of sample heterogeneity and isoform detection, we have taken advantage of 786 GTEx samples (heart: atrial appendage and left ventricle; arteria: aorta and coronary artery) collected from 352 adults ^20^ and 180 iPSC-derived cardiovascular precursor cell (iPSC-CVPC) samples generated from 139 iPSCORE individuals ^18^. We and others previously have shown that iPSC-CVPC display fetal-like gene expression and epigenetic signatures ^21,22^ and that they are composed of varying proportions of fetal-like cardiomyocytes and epicardial-derived cells (EPDCs) including fibroblasts, vascular smooth muscle cells and endothelial cells ^18,23^. Analyzing these 966 bulk RNA-seq samples representing CVS tissues at two developmental stages (fetal and adult) and three distinct types (iPSC-CVPC, heart and arteria), we found that the vast majority of RBPs are expressed at higher levels in the fetal-like iPSC-CVPC compared with adult CVS tissues and play a larger role regulating the expression of fetal-specific isoforms than adult-specific isoforms. We also show that fetal-specific isoforms are more likely to have canonical sequences at their splice sites and to encode proteins that have more functions than their cognate adult-specific isoforms. To examine cell type-specific gene expression differences between fetal-like and adult CVS tissues, we performed cell type deconvolution of the 966 bulk RNA-seq samples. We show that the expression of thousands of genes and isoforms are associated with both CVS developmental stage and cell types (i.e., differentially expressed in the same cell type at different developmental stages), and that developmental stage- and cell type-specific RBP expression drives the observed developmental stage- and cell type-specific isoform usage.

To investigate if the transcriptome during heart failure reverts to a fetal-like program, we compared heart failure samples ^24^ with fetal-like iPSC-CVPC and adult heart samples. We show that while the cellular composition of heart failure and healthy heart samples are similar, RBP genes as a class are overexpressed in heart failure, and more importantly, that the RBPs that are overexpressed substantially overlap but are not exactly the same as those expressed at high levels in fetal-like iPSC-CVPC. Furthermore, we show that overexpression of RBPs in heart failure is associated with a transcriptome-wide isoform switch, which results in the expression of 1,523 fetal heart-specific isoforms.

## Results

### RNA binding proteins show extensive expression differences between fetal-like and adult CVS tissues

To examine global transcriptome-wide gene expression differences between fetal-like iPSC-CVPC and adult CVS tissues we performed dimensionality reduction and clustering of the 966 bulk RNA-seq samples (Figure S1, Table S1, Table S2). We observed that samples divided into three clusters corresponding to iPSC-CVPC, adult heart (atrial appendage and left ventricle samples) and adult arteria (aorta and coronary artery samples, Figure 1A,B, Figure S2A-E, Figure S3, Figure S4, Table S2). For all downstream analyses, we combined atrium and ventricle into “adult heart” and aorta and coronary artery into “adult arteria” to increase statistical power. Cardiomyocytes divide at high rates during fetal development and early childhood, then cell division decreases gradually to <1% per year in adulthood ^25^. To examine cell division rates in iPSC-CVPC compared with the two adult tissues, we calculated G2M and S-phase scores based on the expression of canonical cell cycle markers ^26,27^ and found that iPSC-CVPC were dividing at a significantly higher rate (G2M and S-scores respectively, heart: p = 3.1 × 10^−83^, p = 1.2 × 10^−84^, arteria: p = 1.7 × 10^−79^, p = 1.6 × 10^−79^, Mann-Whitney U test, Figure S2F-G, Table S2). Then we performed pseudotime analysis ^28^ and observed that iPSC-CVPC were significantly at an earlier developmental stage than the two adult CVS tissues (heart: p = 7.3 × 10^−85^, arteria: p = 1.5 × 10^−79^, Mann-Whitney U test, Figure S2H, Table S2). These analyses support the findings of earlier studies ^18,21^ showing that iPSC-CVPC most closely resembles fetal-like CVS tissue.

**Figure 1:**
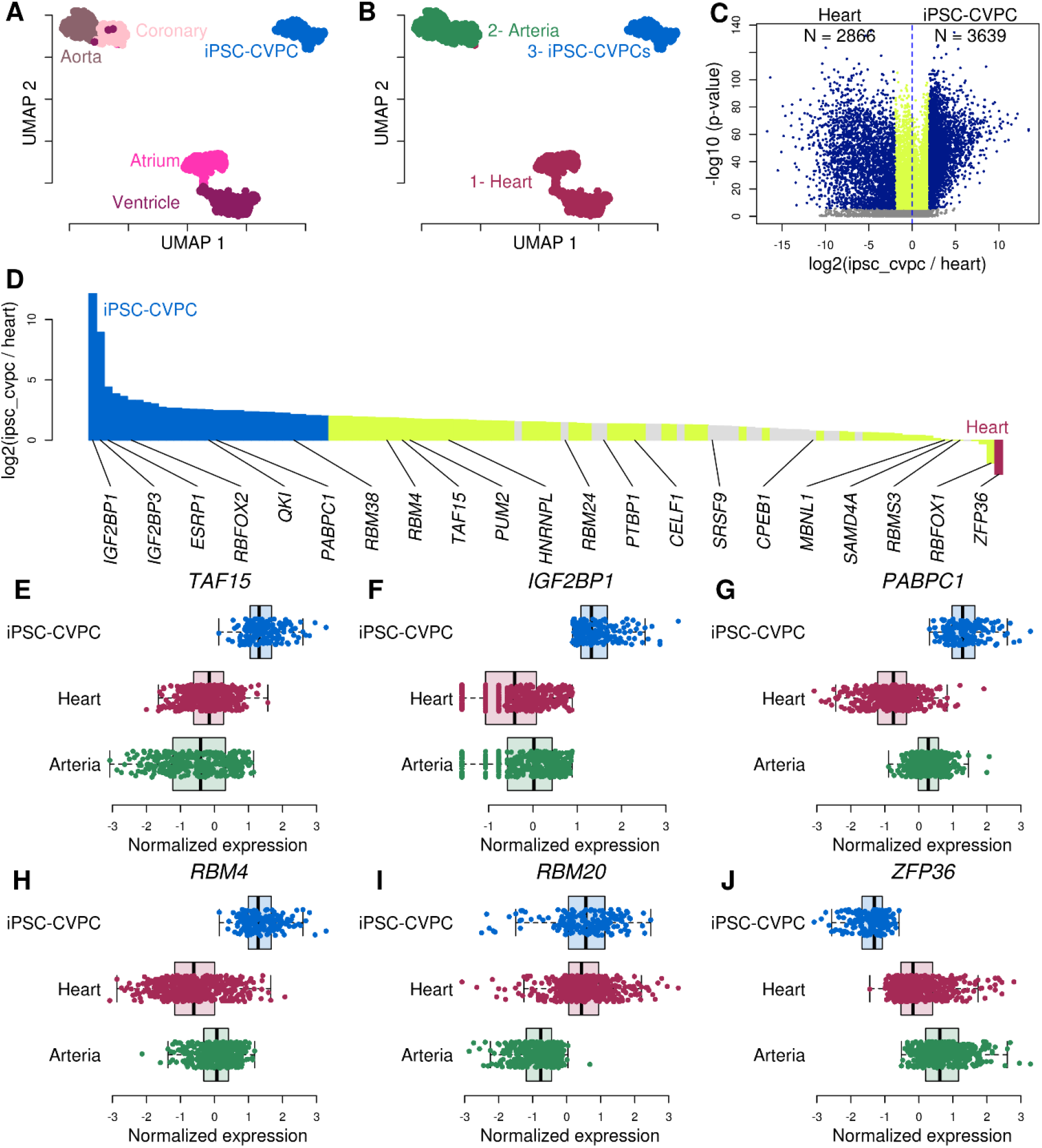
RBPs are overexpressed in fetal-like CVS tissue. (A, B) UMAP plots showing how the 966 RNA-seq samples cluster at varying resolutions: (A) shows the samples according to their tissue of origin (iPSC-CVPC, adult atrium, ventricle, aorta or coronary artery); In panel (B) samples are colored based on clustering at resolution = 0.01. Figures S2A-E and Figure S3 show clustering at different resolutions. (C) Volcano plot showing differential gene expression between iPSC-CVPC and adult heart. The 6,505 genes with greater than four-fold difference (log_2_ ratio > 2 or < −2) are shown in blue (FDR < 0.05) which include 2,866 genes with log_2_ ratio < −2 (adult heart-specific) and 3,639 genes with log_2_ ratio > 2 (iPSC-CVPC-specific). Genes that are significantly differentially expressed (FDR < 0.05) with log_2_ ratio between −2 and 2 are in yellow. Genes that are not differentially expressed (FDR > 0.05) are shown in gray. Volcano plots showing the differential expression between iPSC-CVPC and arteria, and between adult heart and arteria are shown in Figure S5A-B. (D) Barplot showing the log_2_ ratio between the mean expression in iPSC-CVPC and adult heart for 122 RBPs with a known binding motif. iPSC-CVPC-specific RBPs are shown in blue, adult heart-specific RBPs are shown in red. All other differentially expressed RBPs (FDR < 0.05) are shown in yellow. The barplots showing the comparisons between iPSC-CVPC and adult arteria and between adult heart and arteria are shown in Figure S5C-D. The gene symbols of selected RBPs with cardiac- or development-specific functions are shown. (E-J) Boxplot showing gene expression levels (normalized TPM) in iPSC-CVPC, adult heart and adult arteria for six differentially expressed RBPs with cardiac functions. Other RBPs are shown in Figure S6.

We next conducted pairwise gene expression comparisons of the three CVS tissues using linear regression and found 6,505 tissue-specific genes (31.9% of all expressed genes, absolute value of log_2_ ratio > 2 and p-value < 0.05, F-test, after false discovery rate (FDR) correction using Bonferroni’s method) between iPSC-CVPC and adult heart; 5,818 genes (28.5%) between iPSC-CVPC and adult arteria; and 3,358 genes (16.5%) between adult heart and adult arteria (Figure 1C, Figure S5A,B, Table S3). Functional enrichment analysis showed that genes overexpressed in both the adult heart and iPSC-CVPC compared with adult arteria were associated with cardiac muscle function, and that adult arteria-specific genes were enriched for immune response and processes associated with extracellular matrix organization (Table S4). Genes with high expression levels in iPSC-CVPC compared with both adult CVS tissues were most strongly enriched for the RNA binding gene set (GO:0003723, p = 1.8 × 10^−57^ and p = 5.0 × 10^−54^, t-test, compared with adult heart and adult arteria, respectively). This gene set included 122 RNA binding proteins (RBPs) with known binding motifs ^29–31^, of which 93 were overexpressed in iPSC-CVPC (FDR < 0.05) including 33 RBPs that were iPSC-CVPC-specific (log_2_ ratio > 2 and FDR < 0.05), while only four were overexpressed in adult heart (one adult heart-specific: *ZFP36*; log_2_ ratio < −2 and FDR < 0.05) and 24 overexpressed in adult arteria (three adult arteria-specific: *MBNL1*, *RBMS3* and *ZFP36*, Figure 1D, Figure S5C,D). We found that several RBPs known to be involved in fetal cardiac development (including *CELF1*, *HNRNPL*, *RBFOX2* and *RBM24*) and with adult functions (*MBNL1* and *RBFOX1*) ^3–9^ were respectively overexpressed in the iPSC-CVPC and adult CVS tissues (Figure 1D, Figure S6).

We observed that several RBPs associated with embryonic development or cardiac functions were among the most differentially expressed genes between iPSC-CVPC and adult heart. iPSC-CVPC-specific RBPs included: *TAF15*, involved in cell proliferation ^32^ (Figure 1E, Table S3); *IGF2BP1*, which regulates fetal hemoglobin alternative splicing ^33^ (Figure 1F); *PABPC1*, which regulates translation and cardiomyocyte growth ^34^ (Figure 1G); and *RBM4*, which regulates striated muscle differentiation ^35^ (Figure 1H). We also found *RBM20*, which has been shown to have rare inherited variants causing human dilated cardiomyopathy ^36–38^, to be overexpressed in both iPSC-CVPC and adult heart compared with adult arteria (respectively, p = 1.5 × 10^−30^ and p = 1.3 × 10^−55^, Figure 1I), consistent with its predominant expression in striated muscle ^39^ and suggesting that, although it is critical for cardiac function, it likely does not play a role in cardiac development. The only RBP that was significantly overexpressed in both adult heart and adult arteria, compared with fetal-like iPSC-CVPC, was *ZFP36* (heart: log_2_ ratio = −2.8, p = 3.6 × 10^−62^; and arteria: log_2_ ratio = −4.7, p = 4.1 × 10^−78^, Figure 1J). Consistent with its adult-specific expression, this gene is involved in the inhibition of the immune and inflammatory responses highly expressed in atherosclerotic lesions, where it controls inflammatory response by inhibiting the expression of proinflammatory transcripts ^40,41^. These analyses demonstrate that the vast majority of RBPs are expressed at higher levels in the fetal-like iPSC-CVPC than adult CVS tissues, although some RBPs are adult-specific, and suggest that these expression differences contribute to functional cellular differences between the two developmental stages.

### Large-scale isoform switching occurs between fetal-like and adult CVS developmental stages

We hypothesized that the global expression differences of RBPs between developmental stages result in large-scale isoform switching between iPSC-CVPC, adult heart, and adult arteria. Indeed, across all expressed isoforms, we observed 19,270 differentially expressed isoforms between iPSC-CVPC and adult heart, including 1,534 iPSC-CVPC-specific and 3,043 adult heart-specific (Figure 2A, Table S5). Likewise, between iPSC-CVPC and adult arteria, we found 1,785 iPSC-CVPC-specific isoforms and 3,742 adult arteria-specific isoforms, and between adult heart and adult arteria, we found 629 adult heart-specific isoforms and 319 adult arteria-specific isoforms. To confirm that the isoform usage differences between fetal-like iPSC-CVPC and adult heart and arteria were biologically relevant, we tested if we could capture the isoform switching of four cardiac genes known to have well-established developmental stage-specific isoforms (*SCN5A*, *TNNT2*, *ABLIM1* and *TTN*) and for all these genes, we found the expected associations between isoform expression and CVS developmental stage (Figure S7). Interestingly, we discovered that 38 RBPs (31.1% of expressed RBPs) had isoforms with developmental stage-specific expression, including RBPs with known cardiac functions such as *MBNL1*, *FXR1*, *HNRNPM* and *FMR1* (Figure 2B-E).

**Figure 2:**
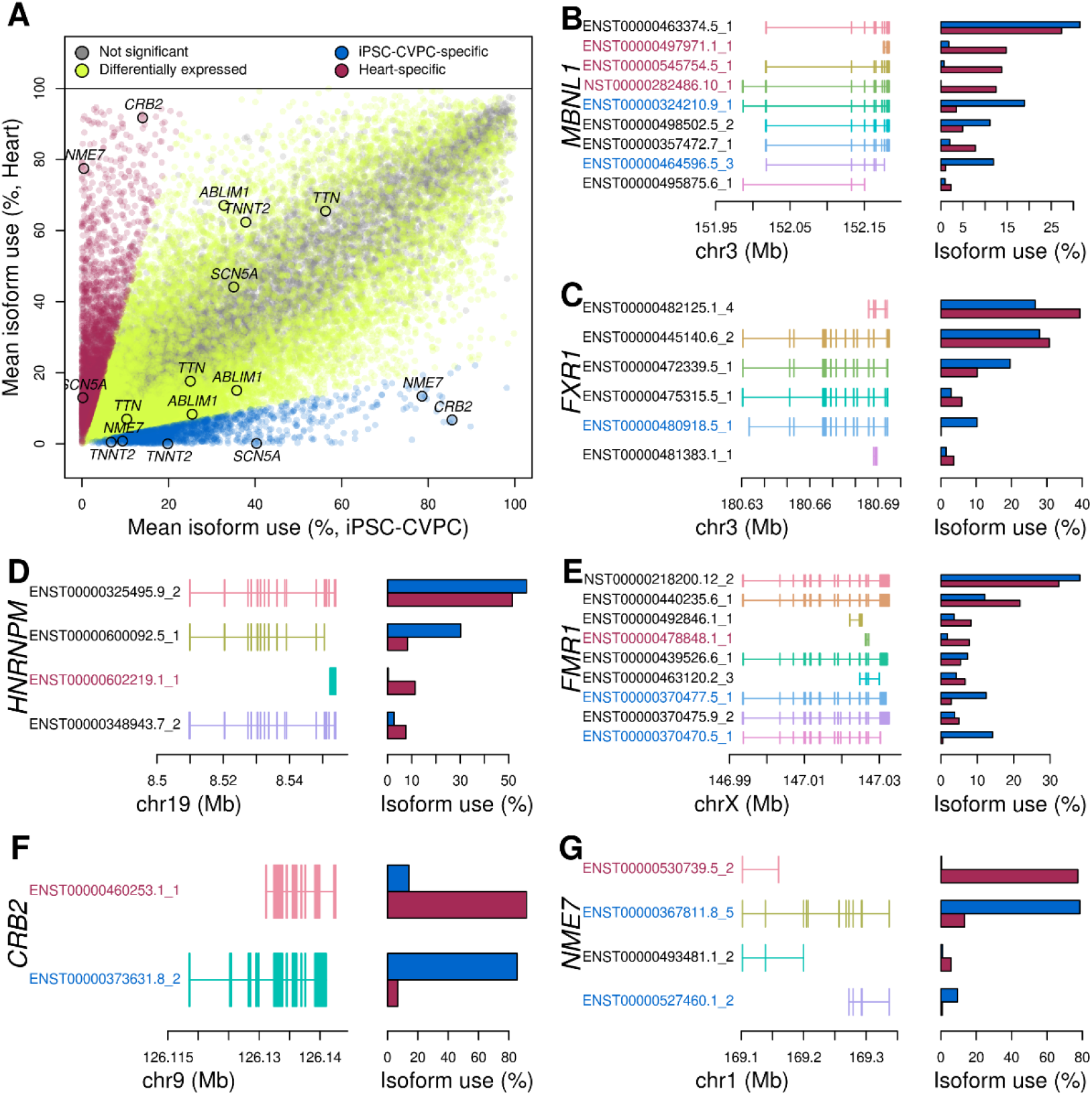
Large-scale isoform switching occurs between fetal-like iPSC-CVPC and heart tissue. (A) Scatter plot showing differential usage between adult heart and iPSC-CVPC for all expressed isoforms. iPSC-CVPC- and adult heart-specific isoforms are shown in blue and purple, respectively (absolute value of the log_2_ ratio between mean isoform use between iPSC-CVPC and adult heart > 2 and FDR < 0.05). All other differentially expressed isoforms are shown in yellow (FDR < 0.05). Isoforms not differentially expressed are shown in gray. Isoform switching occurs between fetal-like iPSC-CVPC and adult heart for four RBPs (B-E) and two genes (F-G) with known cardiac functions. Left side of the figure represents the exon structure of each expressed isoform; the barplots on the right show the mean isoform use (%) of each isoform in iPSC-CVPC (blue) and adult heart (purple). Isoform IDs highlighted in blue are iPSC-CVPC-specific and isoform IDs highlighted in purple are adult heart-specific.

### iPSC-CVPC-specific exons are enriched for functional protein domains

We performed functional enrichment analyses on genes that had at least one differentially expressed isoform between iPSC-CVPC and adult CVS tissues, and found they were enriched for regulating mRNA processing, including alternative splicing and RBP function (Table S6), whereas genes with isoforms differentially expressed between adult heart and adult arteria were enriched for cardiac muscle and mitochondrial functions, which represent physiological differences between the two tissues. To further investigate the differences between CVS developmental stage-specific isoforms, we tested if the iPSC-CVPC-specific and adult heart-specific and arteria-specific isoforms had different associations with functional protein domains. We found that iPSC-CVPC-specific exons were 1.21 more likely to encode for known protein domains ^42^ than adult heart-specific exons (p = 4.4 × 10^−5^, Fisher’s exact test) and 1.15 times than adult arteria-specific exons (p = 1.2 × 10^−3^, Fisher’s exact test, Table S7). We inspected the protein domains of novel developmental stage-specific isoforms in several genes with documented cardiac associated functions. *CRB2,* a gene involved in the differentiation of mesodermal cells ^43^, had one iPSC-CVPC-specific and one shorter adult-associated isoform (Figure 2F) lacking the three C-terminal exons encoding for its extracellular domain, suggesting that its function is likely impaired in adult cardiac tissues. *NME7*, which is involved in centrosome organization and mitotic spindle formation ^44^, had one iPSC-CVPC-specific, one iPSC-CVPC-associated and one adult heart-specific isoform (Figure 2G), which could underlie the significant differences in fetal and adult cardiac cell division rates ^25^. Only the iPSC-CVPC-associated *NME7* isoform (ENST00000367811.8_5) was labeled by Gencode as “protein coding”, whereas the adult heart-specific isoform is a processed transcript which lacks the N-terminal end and is likely not translated into a functional protein. Overall, these findings show that fetal-like iPSC-CVPC-specific isoforms encode proteins that are functionally different from the proteins encoded by their cognate adult-specific isoforms and suggest that adult-specific proteins may have fewer functions than their fetal counterparts.

### iPSC-CVPC-specific exons are enriched for RBP binding and canonical splice sites

To investigate the association between differential isoform usage and differential RBP expression, we tested the overlap of CVS developmental stage-specific isoforms and experimentally determined (eCLIP) binding sites of 39 RBPs ^29^. We found that 33 RBPs were overexpressed in iPSC-CVPC (including 12 iPSC-CVPC-specific) and six were not differentially expressed. We observed that the genomic loci (coding and non-coding sequences) encoding isoforms overexpressed in iPSC-CVPC were more likely to overlap RBP binding sites than the genomic loci encoding isoforms overexpressed in either adult heart or adult arteria, whereas we did not observe significant differences between the two adult tissues (Figure 3A-B, Figure S8, Table S8). We also investigated the occurrence of motifs for the 122 RBPs (GO:0003723) in splice sites of developmental stage-specific exons and found that the splice sites of iPSC-CVPC-specific exons were more likely to harbor RBP motifs than the splice sites of heart-specific and arteria-specific exons (Figure S9, Table S9). Conversely, splice sites for adult heart-specific and arteria-specific exons harbored similar numbers of RBP motifs. These results are consistent with our observation that the vast majority of RBPs are expressed at higher levels in the iPSC-CVPC and indicate that RBPs play a larger role controlling the expression of fetal-specific isoforms than adult-specific isoforms.

**Figure 3:**
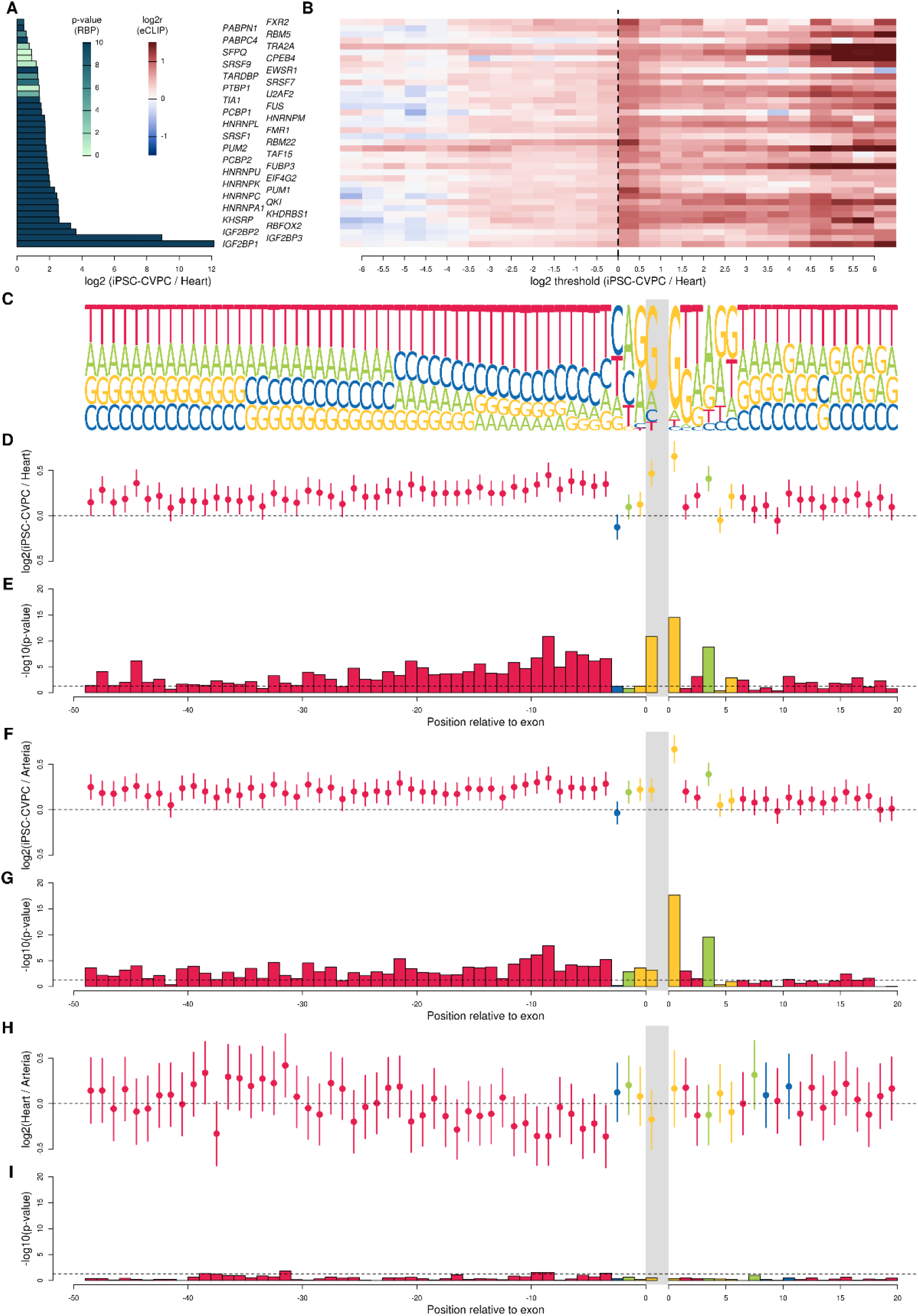
iPSC-CVPC-specific exons enriched for canonical splice sites and polypyrimidine track. (A) Barplot showing the differential expression of RBPs between iPSC-CVPC and adult heart. Colors represent p-values (Table S3). Each row represents one RBP labeled to the right. (B) Heatmap showing enrichment of genomic loci encoding iPSC-CVPC-specific isoforms and adult-specific isoforms for overlapping experimentally determined (eCLIP) RBP binding sites. Each row of the heatmap corresponds to an eCLIP experiment for the indicated RBP sorted as in panel A. Each column corresponds to log_2_ ratio thresholds (mean isoform usage in iPSC-CVPC / mean isoform usage in adult heart; FDR < 0.05). Enrichment was calculated by comparing the proportion of genes with differentially expressed isoforms passing the log_2_ ratio thresholds described on the X axis (Table S5) and overlapping eCLIP peaks against the proportion of genes without any differentially expressed isoforms (FDR > 0.05 for all isoforms) overlapping eCLIP peaks. Genes with iPSC-CVPC-specific isoforms (i.e., positive thresholds) are more likely to overlap eCLIP peaks than genes that have adult heart-specific isoforms (i.e. negative thresholds). (C) Shown is the sequence logo of the 50 bp upstream of the splice acceptor site (left of gray bar) and the 20 bp downstream of the splice donor site (right of gray bar). Gray bar represents the location of the exon. Only the first nucleotide of the exon is shown. Log_2_ ratio between the use of the most common nucleotide at each position between (D) iPSC-CVPC-specific and adult heart-specific exons, (F) iPSC-CVPC-specific and adult arteria-specific exons and (H) adult heart-specific and adult arteria-specific exons. Circles represent the estimate and segments represent the 95% confidence interval as calculated using the *fisher.test* function in R. (E, G, I) Barplots showing the –log_10_ (p-values, Fisher’s exact test) of the enrichments shown in (D, F, H). The Y axis is the same across similar panels: in panel I, three positions have significant differences between adult heart and arteria, with p-values close to 0.05 (dotted line), whereas the majority of comparisons between iPSC-CVPC and each of the two adult tissues are significant in panels E and G. At each position, test results are shown only for the most common nucleotide. Each color represents the tested nucleotide, as shown in Panel C (A = green; C = blue; G = yellow; T = red). Enrichment results for all the nucleotides at all positions are shown in Table S10.

As alternative splicing outcomes are influenced by several factors including splice site strength, we further investigated the splice site sequences of developmental stage-specific exons for differences in the frequency of canonical and non-canonical splice donor and acceptor sites between fetal-like and adult CVS tissues (Figure 3C-I, Table S10). At the splice donor site the first and fourth positions of iPSC-CVPC-specific exons were significantly enriched for the canonical nucleotides (G and A, respectively) compared with both adult heart and adult arteria-specific exons (respectively, p = 3.3 × 10^−15^ and p = 2.1 × 10^−18^ at the first position; p = 1.5 × 10^−9^ and p = 2.7 × 10^−10^ at the fourth position), whereas we did not observe significant differences between the two adult CVS tissues (p = 0.43 and p = 0.46, respectively for the first and fourth position). Since the genomic region immediately upstream of the splice acceptor usually contains a polypyrimidine tract (mostly thymine) of 15-20 nucleotides ^45^, we investigated if iPSC-CVPC-specific exons were enriched for having pyrimidines within the first 100 bp upstream of their splice acceptor site. Of the first 100 bps upstream of the splice acceptor sites, we found that 84 and 81 of the bps for the iPSC-CVPC-specific exons were significantly more likely to be thymine than adult heart-specific exons and adult arteria-specific exons, respectively (FDR < 0.05, Fisher’s exact test, Table S10). Conversely, only eight positions were significantly different between adult heart-specific exons and adult arteria-specific exons. These analyses show that adult-specific exons are more likely to have non-canonical splice sites, consistent with previous studies showing that non-canonical splice sites tend to be tissue-specific and contribute to increasing proteome diversity between adult tissues ^46,47^; on the other hand, iPSC-CVPC-specific exons are more likely to have canonical sequences at their splice sites and the typical upstream polypyrimidine tract, indicating that they are more likely to use conserved splicing machinery.

### Fetal-like and adult CVS transcriptional differences are not due to cellular heterogeneity

We hypothesized that the transcriptional differences between the fetal-like iPSC-CVPC and two adult CVS tissues were due to expression of developmental stage-specific genes and isoforms, while the transcriptional differences between the two adult CVS tissues were a consequence of their cellular composition differences. To test this hypothesis, we deconvoluted and determined the cell type composition ^19^ of each of the 966 bulk RNA-seq samples. Due to the sparsity of human scRNA-seq data, we leveraged the *Tabula Muris* resource ^48^, which contains deep scRNA-seq data from mouse heart and aorta that enables the accurate estimation of cell type proportions in human bulk RNA-seq samples ^19^. The expression levels of marker genes for eight cardiac cell types (Figure S10) was used as input into CIBERSORT ^49^ to estimate cell type composition (Table S11, Table S12). The most common cell type in iPSC-CVPC and adult heart samples was cardiac muscle, whereas adult arteria samples were mainly composed of smooth muscle (Figure 4A, Figure S11). We conducted a linear regression analysis between cell type proportions and gene expression levels (Figure 4A, Figure S12, Table S13) and validated the CIBERSORT-estimated cell type proportions using two methods: 1) by performing gene set enrichment analysis (GSEA) on the effect sizes of each cell type proportion across all genes, we showed that the most significantly enriched gene sets corresponded to the main function associated with each cell type (Table S14); and 2) we observed a strong positive correlation between the effect sizes for each gene and its expression in each cell type from two scRNA-seq CVS studies ^18,50^ (Figure S13).

**Figure 4:**
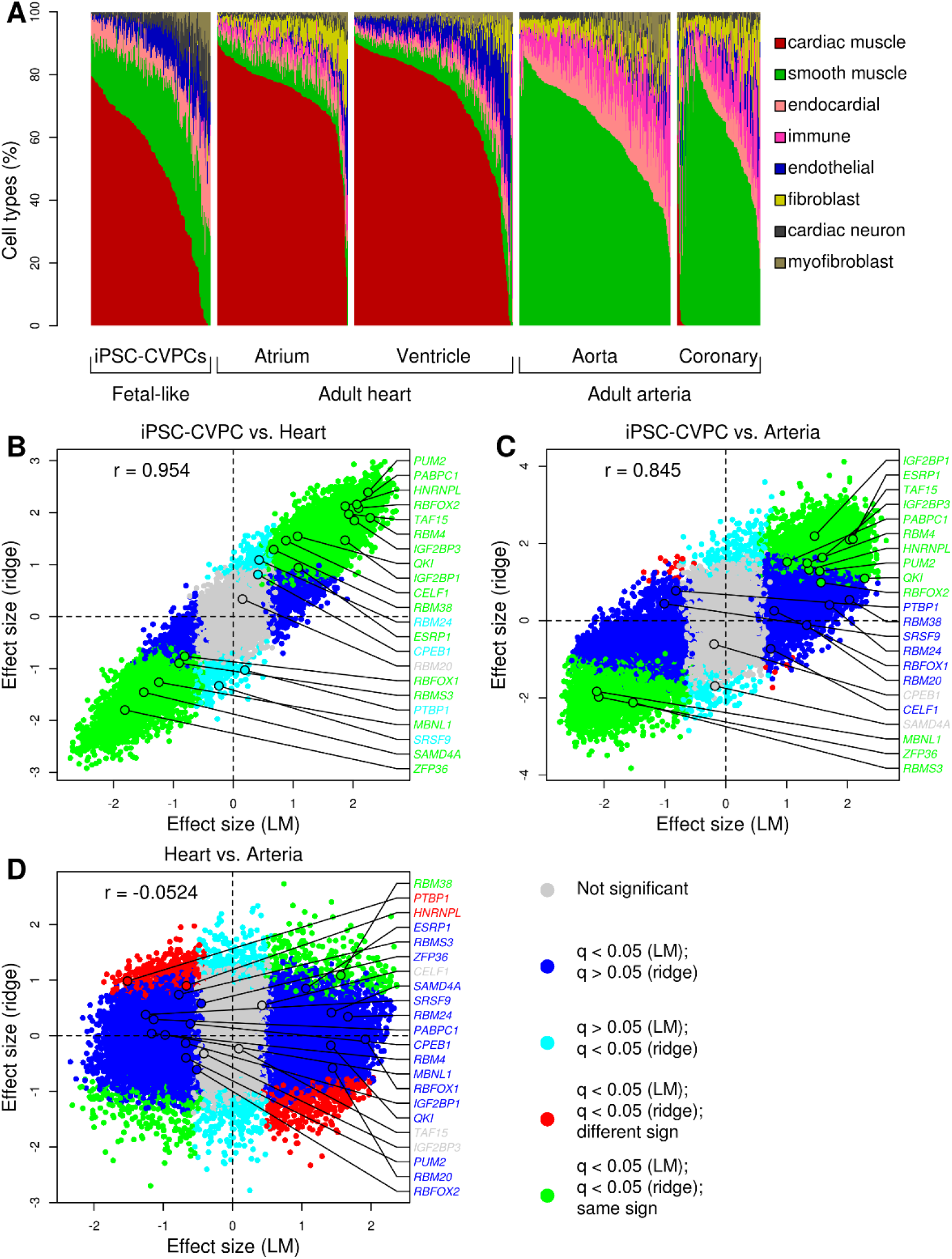
Influence of cell composition on differential expression between CVS tissues. (A) Estimated proportions of cell types across the 966 bulk RNA-seq samples. (B-D) Scatterplots showing the differential gene expression effect size considering cell type proportions as covariates (ridge regression, Y axis) versus without considering cell types (linear model, LM, X axis) for: (B) iPSC-CVPC versus adult heart; (C) iPSC-CVPC versus adult arteria; and (D) adult heart versus adult arteria. Genes differentially expressed in both analyses and have the same direction of effect are shown in green while genes with opposite direction of effect are in red; genes differentially expressed only using the LM are shown in blue; and genes that are differentially expressed only using ridge regression are in cyan. The gene symbols shown to the right of each plot are RBPs with known cardiac or developmental functions shown in Figure 1D.

To test if the expression differences between the CVS tissues were influenced by the different cell type proportions, we repeated the differential expression analysis with cell type proportions as covariates using ridge regression to account for the fact that these covariates are not independent. Between fetal-like iPSC-CVPC and both adult tissues, we found a very strong positive correlation between the effect size of the linear model (which does not take cell type proportions into consideration, Table S3) and the effect size of the ridge regression (r = 0.954 and r = 0.845, respectively, Figure 4B-C, Table S15) indicating that cell type composition only has a minimal effect on the gene expression differences between fetal-like and adult CVS tissues. Of the 6,505 tissue-specific genes between iPSC-CVPC and adult heart and the 5,818 tissue-specific genes between iPSC-CVPC and adult arteria (Table S3), 5,919 (91.0%) and 3,429 (58.9%) were respectively differentially expressed when adjusting for the cell type proportions. Conversely, adult heart and adult arteria were negatively correlated (r = −0.0524) and only 218 (6.5%) heart- or arteria-specific genes identified using the linear model (3,358 genes) were differentially expressed when taking cell type into consideration (Figure 4D). We also tested if differential isoform expression between the three CVS tissues is influenced by the cell type proportions, and observed trends similar to gene expression, with the differences between iPSC-CVPC and the adult tissues being less influenced by cell types than those between adult heart and adult arteria (Figure S14, Table S15). These analyses suggest that fetal-like iPSC-CVPC and adult CVS tissues have significantly different transcriptomes due to the expression of developmental stage-specific genes and isoforms, whereas the transcriptomic differences between the two adult CVS tissues are largely due to different cellular compositions.

### Expression of thousands of genes and isoforms are associated with both CVS developmental stage and cell type

To examine cell-type-specific gene expression differences between fetal-like iPSC-CVPC and the adult CVS tissues, we determined the number of genes whose expression was associated with both CVS developmental stage and cell type. We found 4,636 genes, including 41 RBPs, and 1,356 isoforms that were both differentially expressed between iPSC-CVPC and adult heart (FDR < 0.05) and positively associated with at least one cell type (effect size > 0 and FDR < 0.05, Figure 5A,B, Figure S15, Table S15). Among the RBPs associated with both CVS stage and cell type, several are noteworthy for having functions associated with specific cell types and/or involved in embryonic development (Figure S16). Among the associated isoforms, we identified two of *EXOC7* which differ only by the inclusion/exclusion of exon 7. The isoform expressing exon 7 (ENST00000589210.6_3) was overexpressed in iPSC-CVPC (p = 3.7 × 10^−50^) and positively associated with cardiac muscle proportion (p = 2.1 × 10^−29^), whereas the isoform that does not express exon 7 (ENST00000332065.9_2) was overexpressed in the adult heart (p = 3.7 × 10^−40^), negatively associated with the cardiac muscle proportion (p = 1.5 × 10^−23^) and positively associated with cardiac neuron (p = 6.4 × 10^−30^, Figure 5C-F, Table S15). Polypyrimidine tract-binding protein 1 (PTBP1) has been shown to inhibit exon 7 inclusion most likely by binding to the acceptor site, which leads to exon skipping ^51^. Here, we show that *PTBP1* has the same positive association (p = 5.9 × 10^−13^) with adult cardiac neuron and is similarly expressed at higher levels in the adult heart compared with iPSC-CVPC (p = 3.8 × 10^−18^) as the EXOC7 isoform that does not express exon 7 (ENST00000332065.9_2, Figure 5G). We observed fewer associations that were associated with both CVS developmental stage and cell type between iPSC-CVPC and adult arteria (2,096 genes and 278 isoforms), which suggests that cardiac muscle likely has greater tissue-specific alternative splicing compared with arteria smooth muscle, consistent with previous studies showing that alternative splicing regulation is more prominent in striated muscle and brain than other tissues ^52,53^. Overall, this analysis showed that the expression of thousands of genes and isoforms is associated with both CVS developmental stage and cell types and, more importantly, that RBP developmental stage- and cell type-specific expression drives developmental stage- and cell type-specific usage of isoforms.

**Figure 5:**
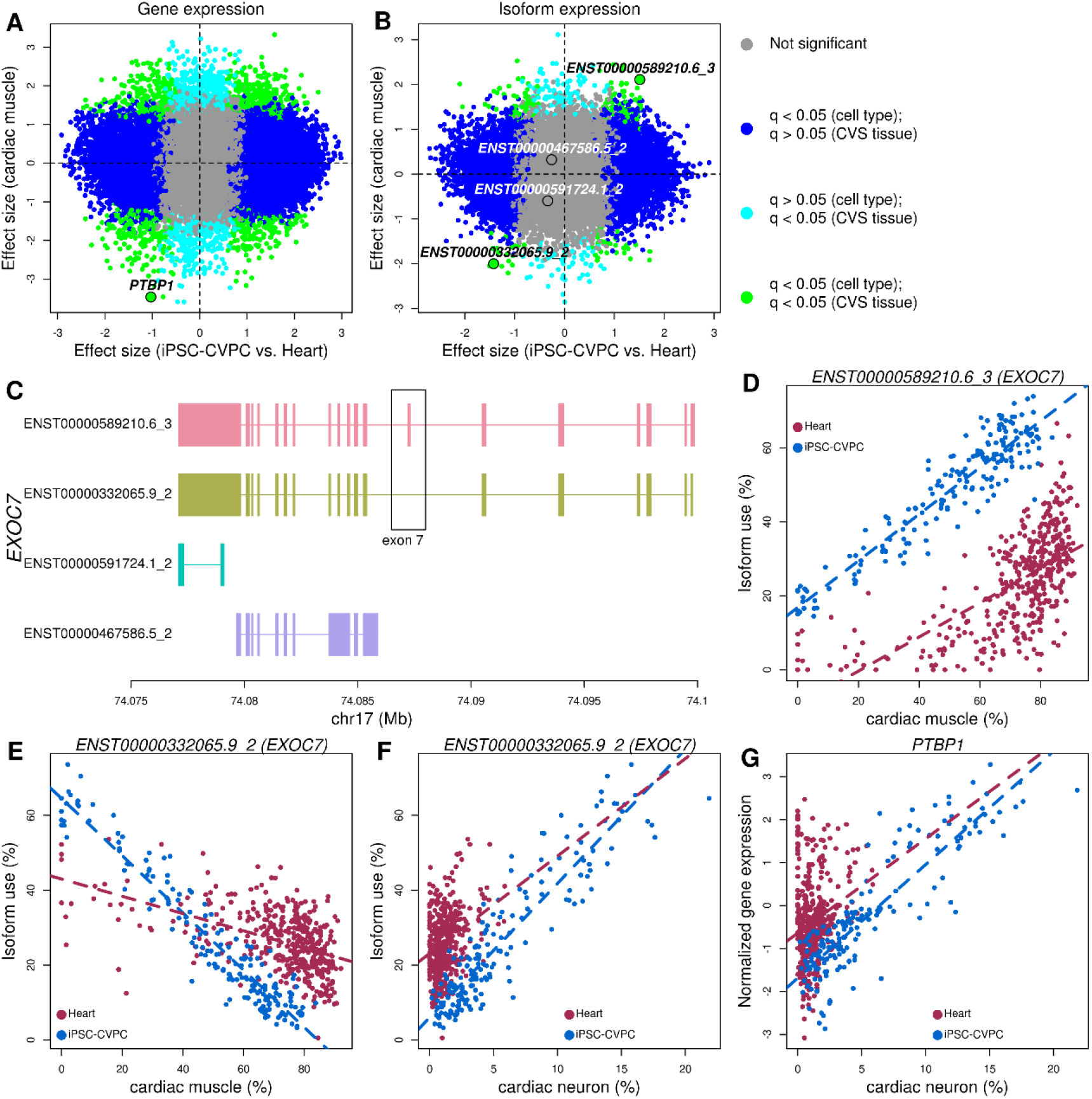
Expression of thousands of genes and isoforms are associated with CVS tissue and cell type. (A,B) Scatterplots showing differential (A) gene and (B) isoform expression effect size between iPSC-CVPC and adult heart (X axis) and the effect size of the association between expression and proportion of cardiac muscle (Y axis). Genes and isoforms significantly associated with both CVS tissue and cardiac muscle proportion are shown in green; genes and isoforms associated only with cell type are in blue; genes and isoforms that are associated only with CVS tissue are in cyan; genes and isoforms not associated with either CVS tissue or cell type proportion are shown in gray. *PTBP1* is shown in (A) and the four *EXOC7* isoforms are shown in (B). (C) Exon structure of the isoforms of *EXOC7*. The two isoforms (ENST00000589210.6_3 and ENST00000332065.9_2) differentially expressed between iPSC-CVPC and adult heart differ in the presence or absence of exon 7. (D-F) Scatterplots showing the association between isoform use (%) and cell type proportion for (D,E) cardiac muscle or (F) cardiac neuron (X axis) in iPSC-CVPC and adult heart (Y axis) for the two most commonly expressed isoforms of *EXOC7*. Dashed lines represent regression lines calculated on each CVS tissue. (G) Scatterplot showing the association between the normalized expression of *PTBP1* and the proportion of cardiac neuron (X axis) in iPSC-CVPC and adult heart. Dashed lines represent regression lines calculated on each CVS tissue.

### Heart failure: reversion to fetal-like RBP and isoform transcriptome

To determine if the CVS transcriptome during heart failure reverts to a fetal-like program ^10,11^, we investigated if RBP expression differences between heart failure samples pre- and post- implantation of mechanical support with a left ventricular assist device (LVAD, Table S16) ^24^ resemble those between fetal-like iPSC-CVPC and adult heart. We first deconvoluted the cell types of 15 paired pre- and post-LVAD samples using CIBERSORT ^49^ and found that their cellular composition was overall comparable to the healthy adult GTEx heart samples (Figure 6A), except for smooth muscle (11.6% in post-LVAD samples compared with 6.8% in healthy adult heart, p = 4.2 × 10^−4^) and other minor cellular differences (Table S17). We then integrated the iPSC-CVPC and GTEx adult heart samples with the 15 paired pre- and post-LVAD samples and performed a principal component analysis using the expression levels of the 122 RBPs. We observed that the pre-LVAD heart failure samples were significantly closer to iPSC-CVPC than post-LVAD (p = 1.3 × 10^−6^, paired t-test), whereas post-LVAD samples were significantly closer to the healthy adult heart samples (p = 2.7 × 10^−8^, paired t-test, Figure 6B,C). To further characterize the role of RBPs in heart failure and if their expression reverts to a fetal-like state, we performed a differential expression analysis between the 15 pre-LVAD heart failure samples, iPSC-CVPC and adult heart using ridge regression taking cell type proportions into consideration (Table S18). While four RBPs were overexpressed and nine were downregulated in heart failure compared with iPSC-CVPC, the mean effect size across all 122 RBPs was not different from zero (p = 0.273, t-test, Figure 6D-E) and hence, overall RBP expression was similar between the two tissues. Conversely, RBPs were expressed at higher levels in heart failure compared with healthy adult heart (p = 0.00128, t-test, Figure 6D,F) in a manner similar to their overexpression in iPSC-CVPC compared with heathy adult heart (p = 2.02 × 10^−13^, t-test, Figure 6D,G). Of note, the 14 RBPs significantly differentially expressed between heart failure and healthy adult heart were also differentially expressed between iPSC-CVPC and healthy adult heart, although three had opposite directions of effect: *HNRNPL*, which is involved in myoblast differentiation ^7^, was overexpressed in iPSC-CVPC but downregulated in heart failure; *MBNL1*, which regulates *SCN5A* alternative splicing and is involved in myotonic dystrophy ^54^, and *SAMD4A*, which is also involved in myotonic dystrophy type 1 ^55^, were downregulated in iPSC-CVPC but overexpressed in heart failure. Interestingly, *QKI*, a gene involved in the inhibition of ischemia/reperfusion-induced apoptosis in cardiac muscle cells and involved in the regulation of the transition of fibroblasts toward profibrotic myofibroblasts in cardiac disease ^56,57^, was the most overexpressed RBP in both iPSC-CVPC and heart failure, compared with healthy adult heart (p = 1.7 × 10^−94^ and p = 6.1 × 10^−13^, respectively, Table S18). These results show that overall RBP function is increased in heart failure, and that the RBPs that are overexpressed strongly overlap but are not exactly the same as those expressed at high levels in fetal-like iPSC-CVPC.

Since we observed a strong overlap of RBP activity in fetal-like iPSC-CVPC and heart failure samples, we reasoned that the transcriptome-wide isoform switching that occurs between iPSC-CVPC and healthy adult heart (Figure 2A) may also occur between healthy heart and heart failure. To test this hypothesis, we performed a differential isoform expression analysis between heart failure, healthy adult heart and iPSC-CVPC. We observed 3,268 differentially expressed isoforms between heart failure samples and adult healthy heart (Table S18), of which 1,523 (46.6%) were differentially expressed with the same trend as when comparing isoform usage between iPSC-CVPC and healthy adult heart, a number 1.85 times higher than expected considering that 10,616 isoforms (37.1% of all tested isoforms) were differentially expressed between iPSC-CVPC and healthy adult heart (odds-ratio = 1.85, p = 1.6 × 10^−56^, Fisher’s exact test, Figure 6H). Interestingly, isoforms for 20 different RBPs were among those that reverted in heart failure to the fetal-like iPSC-CVPC expression pattern (Table S18). Furthermore, the effect sizes of the differential isoform usage between heart failure and healthy adult heart were strongly correlated with the effect sizes of the differential isoform usage between fetal-like iPSC-CVPC and healthy adult heart (r = 0.266, p ≈ 0), indicating that in heart failure a transcriptome-wide isoform switch occurs, which results in the expression of thousands of fetal heart-specific isoforms.

**Figure 6:**
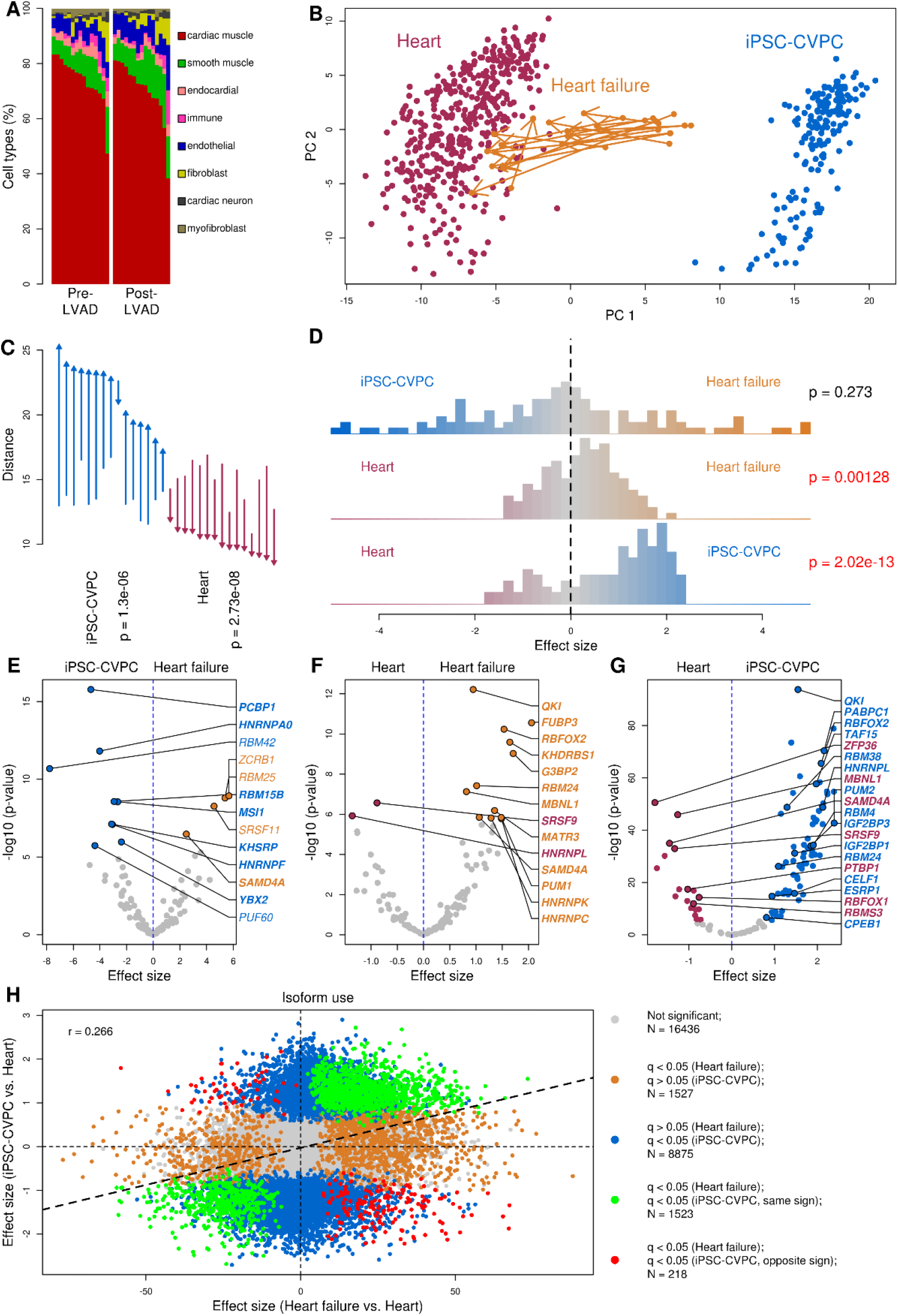
Return to fetal-like expression of RBPs and isoforms in heart failure. (A) Barplot showing the cell type proportions in the 15 heart failure samples pre-LVAD (left) and post-LVAD (right). (B) Scatterplot showing the top two principal components (PC1: 46.1% variance explained; PC2: 20.5% variance explained) calculated on the expression of 122 RBPs in the integrated analysis of iPSC-CVPC, adult heart and heart failure samples. Arrows show the transcriptome changes from pre- to post-LVAD for all heart failure samples. (C) Differences in the average distance between iPSC-CVPC, adult heart and each of the 15 pairs of pre- and post-LVAD heart failure samples calculated on the expression of 122 RBPs shown in B. Arrows indicate average distance and direction (e.g. arrows point up if distance increases) from pre-LVAD to post-LVAD to indicated tissue type. Distance was calculated on the top 10 principal components (87.2% total variance explained). P-values were calculated using paired t-test. (D) Histograms showing the differential expression (effect size) of the 122 RBPs calculated between each pair of CVS tissues (iPSC-CVPC, healthy adult heart and pre-LVAD heart failure). P-values were calculated using one-sample t-test in R (*t.test*, with parameter *mu = 0*). (E-G) Volcano plots showing the differential expression (effect size on the X axis; −log_10_ p-value on the Y axis) between (E) pre-LVAD heart failure and iPSC-CVPC; (F) pre-LVAD heart failure and healthy adult heart; and (G) iPSC-CVPC and adult heart; in (E-G) all differentially expressed RBPs (FDR < 0.05) are shown. In all three panels, the RBPs that are differentially expressed between iPSC-CVPC and healthy adult heart are shown in bold. (H) Scatterplot showing the correlation of the differential isoform usage expression between pre-LVAD heart failure and healthy adult heart (X axis) and between iPSC-CVPC and healthy adult heart (Y axis).

## Discussion

For the past two decades, it has been hypothesized that in heart failure the adult heart reverts to a fetal-like metabolic state and oxygen consumption ^10,11,58^, resulting in the reactivation of the expression of several fetal-specific genes ^12–15^. However, it is currently debated whether these expression changes are largely due to differences in cell type proportions between healthy and diseased hearts ^59,60^, as heart failure triggers the activation of fibroblasts, which ultimately results in cardiac remodeling and fibrosis ^61^. Using the expression levels of marker genes for eight cell types obtained from single cell transcriptomic data, we were able to deconvolute cell type proportions in 996 bulk RNA-seq samples for heart failure, fetal-like iPSC-CVPC, healthy adult heart and arteria ^19,49^. We show that heart failure samples have cell type proportions highly similar to healthy adult heart, with the main difference being a slightly higher fraction of smooth muscle cells in post-LVAD heart failure samples; however, we cannot exclude that this cellular difference may be due to sample collection, rather than reflecting a biological signal, since the heart failure samples were obtained from a different study ^24^. When taking cellular composition into account, we observed that the expression of thousands of genes and isoforms is associated with both CVS developmental stage (fetal-like iPSC-CVPC vs. adult heart) and cell type, which reflects that fact that CVS cell types have developmental stage-specific expression patterns. We also compared the transcriptomes of heart failure samples with both fetal-like iPSC-CVPC and healthy adult heart and observed that heart failure samples have several characteristics that resemble the fetal heart, including the expression of 1,523 of fetal-specific isoforms. Our study indicates that transcriptome-wide isoform switching occurs between fetal and adult heart at the cellular level, and in heart failure a partial reversion to the fetal-specific isoform expression program occurs.

RBPs are a major driver of the underlying transcriptional changes between fetal, healthy and diseased adult heart. The higher expression of RBPs in iPSC-CVPC results in widespread isoform changes between fetal-like and adult CVS tissues. While previous studies have identified only a relatively small number of genes that undergo isoform switching between fetal and adult heart ^8,9,62–70^, we show that thousands of isoforms are fetal- or adult-specific, indicating that alternative splicing plays an important role in driving the transition from fetal to post-natal and adult heart. Interestingly, fetal- and adult-specific isoforms have distinct characteristics, with fetal-specific isoforms being more likely to encode for known protein domains, to contain RBP binding motifs, and have canonical splice sites. These observations, combined with the higher overall expression of RBPs in fetal-like iPSC-CVPC, suggest that, RBPs are necessary to drive alternative splicing in the fetal heart transcriptome, whereas the alternative splicing in the adult heart transcriptome is likely partially driven by RBPs and other factors.

Our study shows that in heart failure reactivation of RBPs results in the expression of 1,523 fetal-specific isoforms which likely contributes to disease pathophysiology including reversion to a fetal-like metabolic state and oxygen consumption ^10,11,58^. Therefore, RBPs, and in particular the 20 that have isoforms differentially expressed in both iPSC-CVPC and heart failure compared with the healthy adult heart, may have two distinct uses in regenerative medicine: 1) to improve the quality and clinical relevance of iPSC-derived cardiac muscle cells; and 2) as novel therapeutic targets for heart failure. Since iPSC-CVPC are fetal-like ^18,21,22^, downregulating the expression of fetal-specific cardiac muscle RBPs would likely result in more mature adult-like cardiac muscle cells, thereby improving the use of iPSC-CVPC as a model system to study the adult heart *in vitro*. Experiments in primates have shown that transplantation of stem cell-derived cardiac cells into infarcted hearts improves cardiac regeneration but has several side-effects, including arrhythmia and ventricular tachycardia ^71,72^; and clinical trials using stem cell-derived cardiac cells or their derivatives are currently underway ^73–75^. Obtaining enhanced, mature iPSC-CVPC for transplantation would likely improve their success in regenerative cardiac medicine. Furthermore, by pharmacologically targeting fetal-specific cardiac muscle RBPs or their isoforms in heart failure, it may be possible to selectively revert diseased cardiac muscle cells to express a healthy adult transcriptome program. Although drugs specifically targeting cardiac muscle RBP isoforms have not been developed, several small molecules and specific antisense oligonucleotides have been shown to selectively inhibit RBPs aberrantly expressed in cancer, resulting in reduced cell viability, migration and/or invasion ^76–80^. These promising studies suggest that future heart failure therapies may target fetal-specific isoforms of cardiac muscle RBPs to either mature and improve iPSC-CVPC for transplantation or directly revert transcriptomes in diseased cardiac muscle cells into healthy adult expression profiles.

## Methods

### iPSCORE subject information and whole genome sequencing

The 139 iPSCORE subjects ^81^ included members of 27 families (2-9 members/family), 7 monozygotic twin pairs, and 55 singletons (Table S1). During recruitment of iPSCORE subjects subject information (sex, age, family, ethnicity and cardiac diseases) was collected. Recruitment was approved by the Institutional Review Boards of the University of California, San Diego and The Salk Institute (project no. 110776ZF). As previously described ^82^, we generated whole genome sequences of the 139 subjects using DNA isolated from blood (130 subjects) or skin fibroblasts (nine subjects) on the HiSeq4000 (Illumina; 150 bp-paired end), SNVs and small indels were called using GATK ^83^. Genotype information for all iPSCORE individuals is available in dbGaP (phs001325).

### iPSC reprogramming, derivation of iPSC-CVPC and generation of RNA-seq data

As previously described in detail ^81,84,85^, we reprogrammed fibroblast samples from 139 iPSCORE individuals using non-integrative Cytotune Sendai virus (Life Technologies) and the 149 iPSCs (7 subjects had 2 or more clones each) were shown to be pluripotent and to have high genomic integrity with no or low numbers of somatic copy-number variants (CNVs). To generate iPSC-derived cardiovascular progenitors (iPSC-CVPC) we used a small molecule differentiation protocol, which included purification of iPSC-CVPC cultures using lactate ^86^, and harvested iPSC-CVPC at day 25 (D25) differentiation ^18,87^. We obtained RNA-seq (150 bp PE) for all 180 iPSC-CVPC (30 of the 149 iPSCs were differentiated two or more times). For gene expression, we obtained transcript per million bp (TPM) as previously described ^18,82^, with one difference: reads were aligned to 62,492 autosomal genes and their corresponding 229,835 isoforms included in Gencode V.34lift37 ^88^. All iPSCORE iPSC-CVPC RNA-seq samples are available through dbGaP (phs000924).

### GTEx samples

FASTQ files for 786 GTEx RNA-seq (75 bp PE) datasets were downloaded from dpGaP (phs000424) and processed with the pipeline described above for the iPSC-CVPC RNA-seq data. GTEx samples were obtained from post mortem tissues from 352 individuals and included 227 aorta, 125 coronary artery, 196 atrial appendage and 238 left ventricle. To obtain genotype information from the 352 GTEx individuals, we downloaded the VCF file for all SNPs and small indels from WGS (phg001219) from dbGaP (phs000424).

### Integration of iPSCORE and GTEx datasets

We generated one matrix that included the expression levels of 62,492 Gencode V.34lift37 genes and 229,835 isoforms in each of the 966 CVS RNA-seq datasets described above (180 iPSCORE and 786 GTEx). To remove genes expressed at low levels and/or only in a small subset of samples, we only considered genes “expressed” if they had a TPM ≥ 1 in at least 10% of samples. For isoform analysis, only expressed genes with at least two expressed isoforms (TPM ≥ 1 and usage >10% in at least 10% of samples) were used. The main difference between the iPSCORE and GTEx RNA-seq data was the read length (150 and 75 bp paired-end, respectively). We could not use read length as a covariate, as the biological differences between iPSC-CVPC and adult hearts would have been nullified due to perfect correlation between the “read length” variable and “developmental stage”. We therefore identified and removed genes and isoforms whose expression levels were associated with read length. We randomly selected 50 iPSCORE RNA-seq samples and trimmed the 3’ end of their read length to 75 bp (Table S2). We determined gene and isoform expression levels in these 50 samples using the pipeline described above. Next, we tested the expression differences between these 75 bp-read samples and their corresponding 150 bp-read samples using paired t-test with Bonferroni correction for FDR. We identified 299 genes and 8,999 isoforms (corresponding to 949 genes) with significant expression level differences (Table S19). This resulted in a total of 20,393 genes and 38,271 isoforms (corresponding to 10,686 genes) used in the differential expression analysis. We quantile-normalized gene and isoform expression levels using the *normalize.quantiles* (preprocessCore) and *qnorm* functions in R, in order to obtain mean expression = 0 and standard deviation = 1 for each gene and isoform.

### Dimensionality reduction of iPSCORE and GTEx samples

To characterize the similarities between the transcriptomes of iPSC-CVPC, adult heart, and adult arteria, we performed principal component analysis (PCA) of bulk RNA-seq on 966 CVS samples using Seurat ^27^. We created a Seurat object using TPM values as input and employed log-normalization and scaling using default parameters of Seurat. We identified the 2,000 most variable features from variance stabilizing transformation (VST) and performed dimensional reduction to obtain 50 principal components. To understand how samples cluster together, we implemented a graph-based approach to cluster samples at different resolutions of 0, 0.01, 0.2, and 0.7. *Clustree* ^89^ was used to build a clustering tree, and dimensional reduction was visualized using UMAP. The top 50 PCs and UMAP coordinates were extracted from the Seurat object using the *Embeddings* function. To estimate the cell cycle phase scores for G2M and S phase for each cardiac sample, we used the *CellCycleScoring* function in Seurat with the reference G2/M and S phase markers in the *cc.genes* object ^26^.

### Estimation of pseudotime using Monocle

To estimate the time reflected by the different developmental stages of the cardiac tissues, we performed pseudotime analyses using Monocle ^28^. Pseudotime was calculated based on the expression of all expressed genes following the standard workflow for constructing developmental trajectories. Pseudotime trajectory was ordered by rooting time (i.e. pseudotime = 0) in iPSC-CVPC using the *GM_state* function.

### Differential expression analysis between CVS tissue types

The three CVS tissue types (iPSC-CVPC, adult heart, adult arteria) are composed of the same cell types but in different proportions, have significantly different ratios of male:female individuals (p = 3.7 × 10^−7^, Fisher’s exact test, Figure S1A) and the same cell types may have different numbers of mitochondria at different developmental stages. For example, cardiac muscle cells have more mitochondria than other cell types found in cardiac tissues, which results in a larger number of RNA-seq reads mapping to the mitochondria and, in turn, a smaller number of reads mapping to autosomes and sex chromosomes. This would result in lower TPM values for samples with high mitochondrial content. To avoid these biases, we performed differential expression analysis between each pair of tissues (iPSC-CVPC vs. adult heart; iPSC-CVPC vs. adult arteria; and adult heart vs. adult arteria) using a linear model (*lm* function in R):

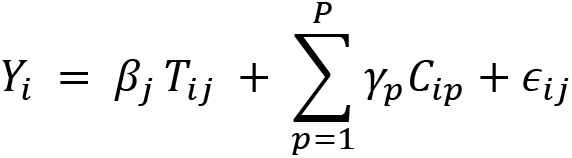

Where *Y*_*i*_ is the phenotype value (normalized gene expression or % isoform use) for sample *i*, *β*_*j*_ is the effect size tissue *j*, *T*_*ij*_ is the tissue type of sample *i*, *C*_*ip*_ is the value of the *p*^th^ covariate for sample *i*, *γ*_*p*_ is the effect size of the *p*^th^ covariate, *P* is the number of covariates used, and ϵ_*ij*_ is the error term for sample *i* at tissue *j*. We used the following covariates: 1) sex; 2) normalized number of RNA-seq reads; 3) % of reads mapping to autosomes or sex chromosome; and 4) % of mitochondrial reads. P-values were corrected using Bonferroni’s method.

### Functional enrichment analysis

Functional enrichment analysis was performed on 12,249 gene sets obtained from MSigDB V.7.1 ^90^. We used gene sets included in the following collections (collection IDs are indicated in parenthesis): Hallmark (h.all); Gene Ontology (GO) biological process (c5.bp), cellular component (c5.cc) and molecular function (c5.mf); canonical pathways included in Biocarta (c2.cp.biocarta), KEGG (c2.cp.kegg) and Reactome (c2.cp.reactome). We performed gene set enrichment analysis (GSEA): for each gene set, we performed a t-test between the effect size (*β*_*j*_in the formulas described above) of each gene included in the gene set and all the expressed genes not included in the gene set. P-values were corrected using Benjamini-Hochberg’s method. We determined the functional enrichment for tissue-associated genes (iPSC-CVPC vs. adult heart; iPSC-CVPC vs. adult arteria; and adult heart vs. adult arteria) and cell type-associated genes.

To determine the functional enrichment of genes with CVS developmental stage-specific isoforms, we obtained all the genes with one or more differentially expressed isoforms (FDR < 0.05 and |log_2_ ratio| > 2) and all the genes that did not have any differentially expressed isoform (FDR > 0.05) and compared their associations with each gene set using Fisher’s exact test. P-values were corrected using Benjamini-Hochberg’s method.

### Defining RBPs

We obtained the position weight matrix (PWM) of 122 RBPs (262 PWMs in total) from three sources: 1) RBPmap V.1.1 ^31^; 2) ENCODE ^29^; and 3) CISBP-RNA V.0.6 ^30^. For all the analyses described in the manuscript, we considered only the 122 RBPs with a known motif obtained from these sources.

### Enrichment of CVS developmental stage-specific isoforms for RBP binding sites

We obtained *in vitro* binding activity of 39 RBPs using enhanced CLIP (eCLIP, 59 experiments) from a recent ENCODE study ^29^. One RBP (*MATR3*) was not expressed in CVS tissues and was removed from further analyses. We downloaded BED files corresponding to biologically reproducible eCLIP peaks and intersected them with the genomic coordinates of each expressed gene. To test whether CVS developmental stage was enriched for having active RBP binding sites, we performed pairwise comparisons (iPSC-CVPC vs. adult heart, iPSC-CVPC vs. adult arteria and adult heart vs. adult arteria) using the following approach:

1. We obtained the list of isoforms overexpressed in stage 1, isoforms overexpressed in stage 2, and converted them to their associated gene bodies. To determine overexpressed isoforms, we used increasingly stringent filtering thresholds, from log_2_ ratio = 0 (the least stringent, including all the isoforms that had a significant p-value after FDR correction in Table S5) to log_2_ ratio > 6. We performed enrichment analysis at each threshold described in Table S8.
2. As background, we used all the genes that did not have any differentially expressed isoforms.
3. We intersected the three lists of gene bodies with the coordinates of eCLIP peaks for each eCLIP experiment.
4. We compared the fraction of gene bodies associated with each stage against background using Fisher’s exact test (*fisher.test* in R).

### Identifying CVS developmental stage-specific exons

To identify exons expressed only in iPSC-CVPC compared with adult heart, we selected all genes that had at least one isoform that was iPSC-CVPC-specific (FDR < 0.05 and log_2_ ratio > 2 in Table S5) and at least one isoform that was adult heart-specific (FDR < 0.05 and log_2_ ratio < −2 in Table S5). Gencode provides unique identifiers for each exon, with exons having the same ID if they belong to different isoforms but have the same coordinates. Therefore we intersected the lists of exons from the iPSC-CVPC-specific and adult heart-specific isoforms and considered as iPSC-CVPC-specific all the exons that we not present in the adult heart-specific isoforms. Likewise, we defined as adult heart-specific exons all the exons that were present only in the adult heart-specific isoform but not in the iPSC-CVPC-specific. We performed similar filtering to obtain CVS stage-specific exons between iPSC-CVPC and adult arteria and between adult heart and adult arteria.

### Enrichment of CVS developmental stage-specific exons for RBP motifs

To determine whether stage-specific exons were enriched with RBP binding motifs, we created a 100-bp window upstream of exons that are specific to iPSC-CVPC, the adult heart or the adult arteria. The first exon of each isoform were excluded. We created a custom motif file, consisting of 262 RBP motifs (corresponding to 122 distinct RBPs) from RBPmap, ENCODE and CISBP-RNA. We tested enrichment using Homer *findMotifsGenome.pl* ^91^ with the custom RBP motif file in a pairwise fashion (iPSC-CVPC vs. adult heart, iPSC-CVPC vs. adult arteria, and adult heart vs. adult arteria). Enrichment was calculated as the log_2_ ratio between the proportion of exons specific for *tissue 1* and those specific for *tissue 2*. Since Homer *findMotifsGenome.pl* tests only for enrichment (but not depletion) of *tissue 1* against background, to determine enrichment in each CVS stage we performed two analyses and then integrated their results: 1) to test the enrichment for *tissue 1*, we used *tissue 2* as background; and 2) to test the enrichment for *tissue 2*, we used *tissue 1* as background. To integrate the two analyses, for each RBP motif, we retained only the analysis with the most significant q-value and adjusted the sign of the log_2_ ratios, with positive values representing enrichment for *tissue 1* and negative values enrichment for *tissue 2*.

### Finding differences in the splice donor and acceptor site sequences

To determine the consensus sequences at the splice donor and acceptor sites of CVS developmental stage-specific exons, we obtained the coordinates of 100 bp upstream and downstream of each exon and converted them to FASTA using *bedtools getfasta*, with options *–s –tab* to obtain strand-specific sequences in a tab-separated text file. We performed pairwise comparison between each pair of CVS tissues (iPSC-CVPC vs. adult heart, iPSC-CVPC vs. adult arteria and adult heart vs. adult arteria). We calculated the frequency of each of the four nucleotides at each position for exons specific for each of the two tested CVS tissues and performed Fisher’s exact tests on each nucleotide to calculate enrichment. In Figure 3D-I, we only show the enrichment values for the most common nucleotide at each position, while Table S10 shows all the tests.

### Finding overlap between CVS developmental stage-specific exons and protein domains

We obtained the mapping on genomic coordinates for >700,000 protein domains from the Prot2HG database ^42^ and intersected their coordinates with the coordinates of the CVS stage-specific exons from each pairwise analysis (iPSC-CVPC vs. adult heart, iPSC-CVPC vs. adult arteria and adult heart vs. adult arteria). To determine enrichment in the fraction of exons that encode for known protein domains between each pair of CVS tissues, we performed Fisher’s exact test.

### Cell composition estimation

To determine the marker genes and their expression levels to use as input for cell type composition deconvolution, we used our recently published pipeline ^19^, which relies on scRNA-seq data from the *Tabula muris* collection ^48^. Although several studies using scRNA-seq in human cardiac tissues have been published, these data are extremely sparse (<50,000 reads/cell) and hence generate less accurate deconvolution results compared with *Tabula muris* (>800,000 reads/cell). We first obtained a Seurat object containing normalized gene expression data from 4,365 FACS-sorted cardiac cells sequenced at high read depth in the *Tabula Muris* collection ^48^. We then used the *FindAllMarkers* function in Seurat with parameters *only.pos = TRUE, min.pct = 0.2, logfc.threshold = 0.1* to determine marker genes in each of the eight cell types defined by *Tabula Muris*: cardiac muscle, cardiac neuron, endocardial, endothelial, fibroblast, immune, myofibroblast and smooth muscle. We found 7,576 genes overexpressed in at least one cell type and combined their expression levels using the Seurat function *AverageExpression* to calculate their average expression in each cell type. To perform cell type deconvolution in all 966 CVS samples, we ran CIBERSORT ^49^ using the average single cell expression levels of all marker genes in each cell type as “signature gene matrix” and their TPM from bulk RNA-seq as “mixture matrix”. We ran CIBERSORT with quantile normalization = FALSE.

### Validation of the estimated cell type proportions

To validate the cell type proportions determined using CIBERSORT, we determined the associations between the expression of each gene and the proportion of each cell type, then we used these associations to: 1) perform GSEA; and 2) test gene expression in two scRNA-seq datasets. To determine the associations between gene expression and each cell type proportion, we used a linear model (*lm* function in R):

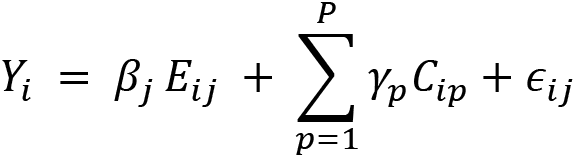

Where *Y*_*i*_ is the phenotype value for sample *i*, *β*_*j*_ is the effect size cell type *j*, *E*_*ij*_ is the proportion of cell type *j* in sample *i*, *C*_*ip*_ is the value of the *p*^th^ covariate for sample *i*, *γ*_*p*_ is the effect size of the *p*^th^ covariate, *P* is the number of covariates used, and *ϵ*_*ij*_ is the error term for sample *i* at cell type *j*. We used the following covariates: 1) sex; 2) normalized number of RNA-seq reads; 3) % of reads mapping to autosomes or sex chromosome; 4) % of mitochondrial reads. P-values were corrected using Bonferroni’s method.

We performed GSEA using the method described above (see section “Functional enrichment analysis”). Briefly, for each gene set, we performed a t-test between the effect size (*β*_*j*_ in the formulas described above) of each gene included in the gene set and all the expressed genes not included in the gene set. P-values were corrected using Benjamini-Hochberg’s method. This analysis shows that each cell type is associated with genes with relevant functions (Table S14).

To further validate the cell type proportions, we used scRNA-seq analysis of 32,026 iPSC-CVPC cells from eight samples ^18^ and 33,050 adult left ventricle cells from 11 samples ^50^. We used CellRanger to aggregate the eight iPSC-CVPC and the 11 adult ventricle samples. Since these two datasets were obtained using different methods from different groups and had different read depths (iPSC-CVPC: 3,030 median UMI/cell and 1,333 expressed genes/cell; adult left ventricle: 463 median UMI/cell and 210 expressed genes/cell post-aggregation), we could not integrate them without confounding real biological differences with batch effects. Therefore, we analyzed them separately to validate the differential expression results that we obtained using bulk RNA-seq.

We aimed at assessing whether the associations between cell type proportion and gene expression could be confirmed using scRNA-seq. To do this, we measured the mean expression in each cell type associated with >100 cells (cardiomyocytes in iPSC-CVPC, cardiac muscle, fibroblasts, endothelium and smooth muscle in the adult heart) and calculated the log_2_ ratio against the mean expression in all the other cells. Mean expression was calculated as *mean(expm1(x)) + 1* in R, where *x* is a vector of expression values across all cells and *expm1* is a function that computes the exponential of a given value minus one. Next, we measured the correlation between the log_2_ ratio calculated in the scRNA-seq samples and the effect size of the association between stage and gene expression in bulk RNA-seq. This correlation was calculated across all genes with at least one read assigned to them in the scRNA-seq and also on the highly expressed genes. Since read depth was different between iPSC-CVPC and adult left ventricle, we used different thresholds: we considered as “highly expressed” all genes expressed in at least 50% of cells for iPSC-CVPC and 10% for adult left ventricle (Figure S13). For all cell types, we observed a significantly positive correlation across all genes and all highly expressed genes, confirming that the associations between gene expression levels and cell type proportions in bulk RNA-seq recapitulate the differences between cell types in scRNA-seq.

To confirm the differences in the stage-specific associations between cardiac muscle proportion and gene expression, we obtained the number of reads mapping to each gene in cardiomyocytes (CM) from iPSC-CVPC and adult cardiac muscle single cells. Next, we normalized the gene expression levels in the two datasets using the *calcNormFactors* function in *edgeR* ^92^ and computed log_2_ ratio between the two stages. We measured the correlation between these log_2_ ratio values and the difference between effect sizes of the association between cardiac muscle proportion and gene expression in bulk RNA-seq for iPSC-CVPC and the association between cardiac muscle proportion and gene expression in bulk RNA-seq for adult heart. The correlation was significantly positive (R = 0.202, p = 4.8 × 10^−162^, Figure S13H), indicating that the CVS developmental stage-specific associations between gene expression and cell type proportions reflect gene expression differences at single-cell resolution.

### Ridge regression analysis to establish associations between expression, CVS stage and cell type proportions

To determine the association between gene expression and isoform usage with CVS stage, taking cell type proportions into consideration, we used ridge regression (*ridge.proj* from the *hdi* package in R), with the following covariates: sex; normalized number of RNA-seq reads; % of reads mapping to autosomes or sex chromosome and % of mitochondrial reads; and the cell type proportions of each of the eight cardiac cell types. P-values were FDR corrected with Bonferroni’s method.

### PCA of heart failure samples with iPSC-CVPC and GTEx samples

To validate the transcriptional changes between adult and fetal gene programs in post-natal pathophysiological conditions, we obtained FASTQ files for bulk RNA-seq of 8 non-ischemic and 7 ischemic heart failing samples before and after mechanical support with left ventricular assist device (i.e. 15 pairs, 30 total samples; GEO ID: GSE46224) ^93^. Reads (100 bp, paired end) were processed with the same pipeline described above for the 966 CVS samples, including the identification of expressed genes and isoforms. All samples had properly paired reads > 98% and percent mRNA reads > 98%. Sample identity check was performed with *plink genome* ^94^ using SNPs with read depth > 10. All 15 sets of paired samples were confirmed as being derived from the same individual (PI_HAT > 98%).

Next, we performed integrative principal component analysis (PCA) on the 30 heart failure samples and the healthy CVS samples (iPSC-CVPC and adult heart) samples using Seurat’s Reference-based Integration Workflow ^27^ where healthy CVS samples were used as reference. We performed integration using TPM values for 121 RBPs (one RBP, *ESRP2*, was not expressed in the heart failure samples). Integration was performed using the standard Seurat workflow (https://satijalab.org/seurat/v3.1/integration.html) on the expression of all 121 RBPs. Principal components were extracted from the integrated object using the Seurat *Embeddings* function.

We calculated pairwise distance between each pair of the 996 samples (966 healthy and 30 heart failure samples) using the *dist* function in R on the top 10 principal components. Next, we obtained the mean distance each of the 30 heart failure samples against iPSC-CVPC, adult heart or adult arteria samples. To determine whether pre- and post-LVAD samples differed in their distance from each of the three cardiac stages, we performed paired t-test.

Differential expression analysis was performed using ridge regression, as described above.

## Data availability

All data used in this manuscript is available through dbGaP, Gene Expression Omnibus (GEO), and Figshare. Bulk RNA-seq and genotype information are available for GTEx and iPSCORE through the dbGaP studies phs000424 (GTEx), phs000924 (iPSCORE, RNA-seq) and phs001325 (iPSCORE whole genome sequencing). scRNA-seq was obtained from dbGaP studies phs000924 (iPSCORE iPSC-CVPC) and phs001539 (adult heart). Heart failure bulk RNA-seq samples were obtained from GEO GSE46224. Supporting data for all the figures and supplemental tables associated with this manuscript have been deposited at Figshare: https://figshare.com/s/b2e70a2ba3a4aac935d9.

## Author information

KAF and MD conceived the study. MD, JPN, TDA and HM performed RNA-seq analysis. HM performed quality check on RNA-seq samples. MD, JPN and HM performed scRNA-seq analysis. MD and TDA performed eCLIP and RBP motif analysis. KAF and MKRD oversaw cellular deconvolution analysis. KAF and ADC oversaw iPSC-CVPC tissue culture and RNA collection for sequencing. MD, JPN and KAF prepared the manuscript.

## Acknowledgements

This work was supported by a California Institute for Regenerative Medicine grant GC1R-06673-B, NSF-CMMI division award 1728497, and NIH grants HG008118, HL107442, DK105541, and DK112155. JPN, TDA and MKRD were supported by T15LM011271. We thank Patrick T. Ellinor and Mark Chaffin for sharing the adult scRNA-seq data prior to release (dbGaP phs001539).

## References

1 Baralle, F. E. & Giudice, J. Alternative splicing as a regulator of development and tissue identity. Nat Rev Mol Cell Biol 18, 437–451, doi:10.1038/nrm.2017.27 (2017).

2 de Bruin, R. G., Rabelink, T. J., van Zonneveld, A. J. & van der Veer, E. P. Emerging roles for RNA-binding proteins as effectors and regulators of cardiovascular disease. Eur Heart J 38, 1380–1388, doi:10.1093/eurheartj/ehw567 (2017).

3 Kalsotra, A. et al. A postnatal switch of CELF and MBNL proteins reprograms alternative splicing in the developing heart. Proc Natl Acad Sci U S A 105, 20333–20338, doi:10.1073/pnas.0809045105 (2008).

4 Gao, C. et al. RBFox1-mediated RNA splicing regulates cardiac hypertrophy and heart failure. J Clin Invest 126, 195–206, doi:10.1172/JCI84015 (2016).

5 Gazzara, M. R. et al. Ancient antagonism between CELF and RBFOX families tunes mRNA splicing outcomes. Genome Res 27, 1360–1370, doi:10.1101/gr.220517.117 (2017).

6 Zhang, M. et al. Rbm24, a target of p53, is necessary for proper expression of p53 and heart development. Cell Death Differ 25, 1118–1130, doi:10.1038/s41418-017-0029-8 (2018).

7 Zhao, Y. et al. MyoD induced enhancer RNA interacts with hnRNPL to activate target gene transcription during myogenic differentiation. Nat Commun 10, 5787, doi:10.1038/s41467-019-13598-0 (2019).

8 Loiselle, J. J. & Sutherland, L. C. Differential downregulation of Rbm5 and Rbm10 during skeletal and cardiac differentiation. In Vitro Cell Dev Biol Anim 50, 331–339, doi:10.1007/s11626-013-9708-z (2014).

9 Giudice, J. et al. Alternative splicing regulates vesicular trafficking genes in cardiomyocytes during postnatal heart development. Nat Commun 5, 3603, doi:10.1038/ncomms4603 (2014).

10 Rajabi, M., Kassiotis, C., Razeghi, P. & Taegtmeyer, H. Return to the fetal gene program protects the stressed heart: a strong hypothesis. Heart Fail Rev 12, 331–343, doi:10.1007/s10741-007-9034-1 (2007).

11 Taegtmeyer, H., Sen, S. & Vela, D. Return to the fetal gene program: a suggested metabolic link to gene expression in the heart. Ann N Y Acad Sci 1188, 191–198, doi:10.1111/j.1749-6632.2009.05100.x (2010).

12 Black, F. M. et al. The vascular smooth muscle alpha-actin gene is reactivated during cardiac hypertrophy provoked by load. J Clin Invest 88, 1581–1588, doi:10.1172/JCI115470 (1991).

13 Kinoshita, H. et al. T-type Ca2+ channel blockade prevents sudden death in mice with heart failure. Circulation 120, 743–752, doi:10.1161/CIRCULATIONAHA.109.857011 (2009).

14 Currie, S. & Smith, G. L. Enhanced phosphorylation of phospholamban and downregulation of sarco/endoplasmic reticulum Ca2+ ATPase type 2 (SERCA 2) in cardiac sarcoplasmic reticulum from rabbits with heart failure. Cardiovasc Res 41, 135–146, doi:10.1016/s0008-6363(98)00241-7 (1999).

15 Carniel, E. et al. Alpha-myosin heavy chain: a sarcomeric gene associated with dilated and hypertrophic phenotypes of cardiomyopathy. Circulation 112, 54–59, doi:10.1161/CIRCULATIONAHA.104.507699 (2005).

16 Xu, X. et al. ASF/SF2-regulated CaMKIIdelta alternative splicing temporally reprograms excitation-contraction coupling in cardiac muscle. Cell 120, 59–72, doi:10.1016/j.cell.2004.11.036 (2005).

17 Beqqali, A. Alternative splicing in cardiomyopathy. Biophys Rev 10, 1061–1071, doi:10.1007/s12551-018-0439-y (2018).

18 D’Antonio-Chronowska, A. et al. Association of Human iPSC Gene Signatures and X Chromosome Dosage with Two Distinct Cardiac Differentiation Trajectories. Stem Cell Reports 13, 924–938, doi:10.1016/j.stemcr.2019.09.011 (2019).

19 Donovan, M. K. R., D’Antonio-Chronowska, A., D’Antonio, M. & Frazer, K. A. Cellular deconvolution of GTEx tissues powers discovery of disease and cell-type associated regulatory variants. Nat Commun 11, 955, doi:10.1038/s41467-020-14561-0 (2020).

20 Consortium, G. T. The GTEx Consortium atlas of genetic regulatory effects across human tissues. Science 369, 1318–1330, doi:10.1126/science.aaz1776 (2020).

21 Benaglio, P. et al. Allele-specific NKX2-5 binding underlies multiple genetic associations with human electrocardiographic traits. Nat Genet 51, 1506–1517, doi:10.1038/s41588-019-0499-3 (2019).

22 Karakikes, I., Ameen, M., Termglinchan, V. & Wu, J. C. Human induced pluripotent stem cell-derived cardiomyocytes: insights into molecular, cellular, and functional phenotypes. Circ Res 117, 80–88, doi:10.1161/CIRCRESAHA.117.305365 (2015).

23 Friedman, C. E. et al. Single-Cell Transcriptomic Analysis of Cardiac Differentiation from Human PSCs Reveals HOPX-Dependent Cardiomyocyte Maturation. Cell Stem Cell 23, 586–598 e588, doi:10.1016/j.stem.2018.09.009 (2018).

24 Yang, K. C. et al. Deep RNA sequencing reveals dynamic regulation of myocardial noncoding RNAs in failing human heart and remodeling with mechanical circulatory support. Circulation 129, 1009–1021, doi:10.1161/CIRCULATIONAHA.113.003863 (2014).

25 Bergmann, O. et al. Dynamics of Cell Generation and Turnover in the Human Heart. Cell 161, 1566–1575, doi:10.1016/j.cell.2015.05.026 (2015).

26 Nestorowa, S. et al. A single-cell resolution map of mouse hematopoietic stem and progenitor cell differentiation. Blood 128, e20–31, doi:10.1182/blood-2016-05-716480 (2016).

27 Butler, A., Hoffman, P., Smibert, P., Papalexi, E. & Satija, R. Integrating single-cell transcriptomic data across different conditions, technologies, and species. Nat Biotechnol 36, 411–420, doi:10.1038/nbt.4096 (2018).

28 Qiu, X. et al. Reversed graph embedding resolves complex single-cell trajectories. Nature methods 14, 979–982, doi:10.1038/nmeth.4402 (2017).

29 Van Nostrand, E. L. et al. A large-scale binding and functional map of human RNA-binding proteins. Nature 583, 711–719, doi:10.1038/s41586-020-2077-3 (2020).

30 Ray, D. et al. A compendium of RNA-binding motifs for decoding gene regulation. Nature 499, 172–177, doi:10.1038/nature12311 (2013).

31 Paz, I., Kosti, I., Ares, M., Jr., Cline, M. & Mandel-Gutfreund, Y. RBPmap: a web server for mapping binding sites of RNA-binding proteins. Nucleic Acids Res 42, W361–367, doi:10.1093/nar/gku406 (2014).

32 Ballarino, M. et al. TAF15 is important for cellular proliferation and regulates the expression of a subset of cell cycle genes through miRNAs. Oncogene 32, 4646–4655, doi:10.1038/onc.2012.490 (2013).

33 de Vasconcellos, J. F. et al. IGF2BP1 overexpression causes fetal-like hemoglobin expression patterns in cultured human adult erythroblasts. Proc Natl Acad Sci U S A 114, E5664–E5672, doi:10.1073/pnas.1609552114 (2017).

34 Chorghade, S. et al. Poly(A) tail length regulates PABPC1 expression to tune translation in the heart. Elife 6, doi:10.7554/eLife.24139 (2017).

35 Lin, J. C. & Tarn, W. Y. RNA-binding motif protein 4 translocates to cytoplasmic granules and suppresses translation via argonaute2 during muscle cell differentiation. J Biol Chem 284, 34658–34665, doi:10.1074/jbc.M109.032946 (2009).

36 van den Hoogenhof, M. M. G. et al. RBM20 Mutations Induce an Arrhythmogenic Dilated Cardiomyopathy Related to Disturbed Calcium Handling. Circulation 138, 1330–1342, doi:10.1161/CIRCULATIONAHA.117.031947 (2018).

37 Brauch, K. M. et al. Mutations in ribonucleic acid binding protein gene cause familial dilated cardiomyopathy. J Am Coll Cardiol 54, 930–941, doi:10.1016/j.jacc.2009.05.038 (2009).

38 Beqqali, A. et al. A mutation in the glutamate-rich region of RNA-binding motif protein 20 causes dilated cardiomyopathy through missplicing of titin and impaired Frank-Starling mechanism. Cardiovasc Res 112, 452–463, doi:10.1093/cvr/cvw192 (2016).

39 Guo, W. et al. RBM20, a gene for hereditary cardiomyopathy, regulates titin splicing. Nat Med 18, 766–773, doi:10.1038/nm.2693 (2012).

40 Zhang, H. et al. mRNA-binding protein ZFP36 is expressed in atherosclerotic lesions and reduces inflammation in aortic endothelial cells. Arterioscler Thromb Vasc Biol 33, 1212–1220, doi:10.1161/ATVBAHA.113.301496 (2013).

41 Moore, M. J. et al. ZFP36 RNA-binding proteins restrain T cell activation and anti-viral immunity. Elife 7, doi:10.7554/eLife.33057 (2018).

42 Stanek, D. et al. Prot2HG: a database of protein domains mapped to the human genome. Database (Oxford) 2020, doi:10.1093/database/baz161 (2020).

43 Ramkumar, N. et al. Crumbs2 promotes cell ingression during the epithelial-to-mesenchymal transition at gastrulation. Nat Cell Biol 18, 1281–1291, doi:10.1038/ncb3442 (2016).

44 Liu, P., Choi, Y. K. & Qi, R. Z. NME7 is a functional component of the gamma-tubulin ring complex. Mol Biol Cell 25, 2017–2025, doi:10.1091/mbc.E13-06-0339 (2014).

45 Wagner, E. J. & Garcia-Blanco, M. A. Polypyrimidine tract binding protein antagonizes exon definition. Mol Cell Biol 21, 3281–3288, doi:10.1128/MCB.21.10.3281-3288.2001 (2001).

46 Parada, G. E., Munita, R., Cerda, C. A. & Gysling, K. A comprehensive survey of non-canonical splice sites in the human transcriptome. Nucleic Acids Res 42, 10564–10578, doi:10.1093/nar/gku744 (2014).

47 Sibley, C. R., Blazquez, L. & Ule, J. Lessons from non-canonical splicing. Nat Rev Genet 17, 407–421, doi:10.1038/nrg.2016.46 (2016).

48 Tabula Muris, C. et al. Single-cell transcriptomics of 20 mouse organs creates a Tabula Muris. Nature 562, 367–372, doi:10.1038/s41586-018-0590-4 (2018).

49 Newman, A. M. et al. Determining cell type abundance and expression from bulk tissues with digital cytometry. Nat Biotechnol 37, 773–782, doi:10.1038/s41587-019-0114-2 (2019).

50 Tucker, N. R. et al. Transcriptional and Cellular Diversity of the Human Heart. Circulation, doi:10.1161/CIRCULATIONAHA.119.045401 (2020).

51 Georgilis, A. et al. PTBP1-Mediated Alternative Splicing Regulates the Inflammatory Secretome and the Pro-tumorigenic Effects of Senescent Cells. Cancer Cell 34, 85–102 e109, doi:10.1016/j.ccell.2018.06.007 (2018).

52 Buljan, M. et al. Tissue-specific splicing of disordered segments that embed binding motifs rewires protein interaction networks. Mol Cell 46, 871–883, doi:10.1016/j.molcel.2012.05.039 (2012).

53 Ellis, J. D. et al. Tissue-specific alternative splicing remodels protein-protein interaction networks. Mol Cell 46, 884–892, doi:10.1016/j.molcel.2012.05.037 (2012).

54 Freyermuth, F. et al. Splicing misregulation of SCN5A contributes to cardiac-conduction delay and heart arrhythmia in myotonic dystrophy. Nat Commun 7, 11067, doi:10.1038/ncomms11067 (2016).

55 de Haro, M. et al. Smaug/SAMD4A restores translational activity of CUGBP1 and suppresses CUG-induced myopathy. PLoS Genet 9, e1003445, doi:10.1371/journal.pgen.1003445 (2013).

56 Chothani, S. et al. Widespread Translational Control of Fibrosis in the Human Heart by RNA-Binding Proteins. Circulation 140, 937–951, doi:10.1161/CIRCULATIONAHA.119.039596 (2019).

57 Guo, W. et al. RNA binding protein QKI inhibits the ischemia/reperfusion-induced apoptosis in neonatal cardiomyocytes. Cell Physiol Biochem 28, 593–602, doi:10.1159/000335755 (2011).

58 Razeghi, P. et al. Metabolic gene expression in fetal and failing human heart. Circulation 104, 2923–2931, doi:10.1161/hc4901.100526 (2001).

59 Hocker, J. D. et al. Cardiac Cell Type-Specific Gene Regulatory Programs and Disease Risk Association. BioRxiV, doi:https://doi.org/10.1101/2020.09.11.291724 (2020).

60 Spurrell, C. H. et al. Genome-Wide Fetalization of Enhancer Architecture in Heart Disease. bioRxiv, doi:https://doi.org/10.1101/591362 (2019).

61 Humeres, C. & Frangogiannis, N. G. Fibroblasts in the Infarcted, Remodeling, and Failing Heart. JACC Basic Transl Sci 4, 449–467, doi:10.1016/j.jacbts.2019.02.006 (2019).

62 Veerman, C. C. et al. Switch From Fetal to Adult SCN5A Isoform in Human Induced Pluripotent Stem Cell-Derived Cardiomyocytes Unmasks the Cellular Phenotype of a Conduction Disease-Causing Mutation. J Am Heart Assoc 6, doi:10.1161/JAHA.116.005135 (2017).

63 Townsend, P. J. et al. Human cardiac troponin T: identification of fetal isoforms and assignment of the TNNT2 locus to chromosome 1q. Genomics 21, 311–316, doi:10.1006/geno.1994.1271 (1994).

64 Gomes, A. V. et al. Cardiac troponin T isoforms affect the Ca(2+) sensitivity of force development in the presence of slow skeletal troponin I: insights into the role of troponin T isoforms in the fetal heart. J Biol Chem 279, 49579–49587, doi:10.1074/jbc.M407340200 (2004).

65 Lahmers, S., Wu, Y., Call, D. R., Labeit, S. & Granzier, H. Developmental control of titin isoform expression and passive stiffness in fetal and neonatal myocardium. Circ Res 94, 505–513, doi:10.1161/01.RES.0000115522.52554.86 (2004).

66 Stevens, J. et al. Analysis of the asymmetrically expressed Ablim1 locus reveals existence of a lateral plate Nodal-independent left sided signal and an early, left-right independent role for nodal flow. BMC Dev Biol 10, 54, doi:10.1186/1471-213X-10-54 (2010).

67 Ohsawa, N., Koebis, M., Mitsuhashi, H., Nishino, I. & Ishiura, S. ABLIM1 splicing is abnormal in skeletal muscle of patients with DM1 and regulated by MBNL, CELF and PTBP1. Genes Cells 20, 121–134, doi:10.1111/gtc.12201 (2015).

68 Freiburg, A. et al. Series of exon-skipping events in the elastic spring region of titin as the structural basis for myofibrillar elastic diversity. Circ Res 86, 1114–1121, doi:10.1161/01.res.86.11.1114 (2000).

69 Neagoe, C. et al. Titin isoform switch in ischemic human heart disease. Circulation 106, 1333–1341, doi:10.1161/01.cir.0000029803.93022.93 (2002).

70 Roof, D. J., Hayes, A., Adamian, M., Chishti, A. H. & Li, T. Molecular characterization of abLIM, a novel actin-binding and double zinc finger protein. J Cell Biol 138, 575–588, doi:10.1083/jcb.138.3.575 (1997).

71 Chong, J. J. et al. Human embryonic-stem-cell-derived cardiomyocytes regenerate non-human primate hearts. Nature 510, 273–277, doi:10.1038/nature13233 (2014).

72 Shiba, Y. et al. Allogeneic transplantation of iPS cell-derived cardiomyocytes regenerates primate hearts. Nature 538, 388–391, doi:10.1038/nature19815 (2016).

73 Menasche, P. et al. Transplantation of Human Embryonic Stem Cell-Derived Cardiovascular Progenitors for Severe Ischemic Left Ventricular Dysfunction. J Am Coll Cardiol 71, 429–438, doi:10.1016/j.jacc.2017.11.047 (2018).

74 Traverse, J. H. et al. First-in-Man Study of a Cardiac Extracellular Matrix Hydrogel in Early and Late Myocardial Infarction Patients. JACC Basic Transl Sci 4, 659–669, doi:10.1016/j.jacbts.2019.07.012 (2019).

75 Borow, K. M., Yaroshinsky, A., Greenberg, B. & Perin, E. C. Phase 3 DREAM-HF Trial of Mesenchymal Precursor Cells in Chronic Heart Failure. Circ Res 125, 265–281, doi:10.1161/CIRCRESAHA.119.314951 (2019).

76 Hong, S. RNA Binding Protein as an Emerging Therapeutic Target for Cancer Prevention and Treatment. J Cancer Prev 22, 203–210, doi:10.15430/JCP.2017.22.4.203 (2017).

77 Graff, J. R. et al. Therapeutic suppression of translation initiation factor eIF4E expression reduces tumor growth without toxicity. J Clin Invest 117, 2638–2648, doi:10.1172/JCI32044 (2007).

78 Muralidharan, R. et al. Tumor-targeted Nanoparticle Delivery of HuR siRNA Inhibits Lung Tumor Growth In Vitro and In Vivo By Disrupting the Oncogenic Activity of the RNA-binding Protein HuR. Mol Cancer Ther 16, 1470–1486, doi:10.1158/1535-7163.MCT-17-0134 (2017).

79 Huang, Y. H. et al. Delivery of Therapeutics Targeting the mRNA-Binding Protein HuR Using 3DNA Nanocarriers Suppresses Ovarian Tumor Growth. Cancer Res 76, 1549–1559, doi:10.1158/0008-5472.CAN-15-2073 (2016).

80 Wu, X. et al. Identification and validation of novel small molecule disruptors of HuR-mRNA interaction. ACS Chem Biol 10, 1476–1484, doi:10.1021/cb500851u (2015).

81 Panopoulos, A. D. et al. iPSCORE: A Resource of 222 iPSC Lines Enabling Functional Characterization of Genetic Variation across a Variety of Cell Types. Stem Cell Reports 8, 1086–1100, doi:10.1016/j.stemcr.2017.03.012 (2017).

82 DeBoever, C. et al. Large-Scale Profiling Reveals the Influence of Genetic Variation on Gene Expression in Human Induced Pluripotent Stem Cells. Cell Stem Cell 20, 533–546 e537, doi:10.1016/j.stem.2017.03.009 (2017).

83 McKenna, A. et al. The Genome Analysis Toolkit: a MapReduce framework for analyzing next-generation DNA sequencing data. Genome Res 20, 1297–1303, doi:10.1101/gr.107524.110 (2010).

84 D’Antonio, M. et al. Insights into the Mutational Burden of Human Induced Pluripotent Stem Cells from an Integrative Multi-Omics Approach. Cell Rep 24, 883–894, doi:10.1016/j.celrep.2018.06.091 (2018).

85 Jakubosky, D. et al. Properties of structural variants and short tandem repeats associated with gene expression and complex traits. Nat Commun 11, 2927, doi:10.1038/s41467-020-16482-4 (2020).

86 Tohyama, S. et al. Distinct metabolic flow enables large-scale purification of mouse and human pluripotent stem cell-derived cardiomyocytes. Cell Stem Cell 12, 127–137, doi:10.1016/j.stem.2012.09.013 (2013).

87 D’Antonio-Chronowska, A., D’Antonio, M. & Frazer, K. A. In vitro Differentiation of Human iPSC-derived Cardiovascular Progenitor Cells (iPSC-CVPCs) Bio-Protocol 10, doi:10.21769/BioProtoc.3755 (2020).

88 Frankish, A. et al. GENCODE reference annotation for the human and mouse genomes. Nucleic Acids Res 47, D766–D773, doi:10.1093/nar/gky955 (2019).

89 Zappia, L. & Oshlack, A. Clustering trees: a visualization for evaluating clusterings at multiple resolutions. Gigascience 7, doi:10.1093/gigascience/giy083 (2018).

90 Subramanian, A. et al. Gene set enrichment analysis: a knowledge-based approach for interpreting genome-wide expression profiles. Proc Natl Acad Sci U S A 102, 15545–15550, doi:10.1073/pnas.0506580102 (2005).

91 Heinz, S. et al. Simple combinations of lineage-determining transcription factors prime cis-regulatory elements required for macrophage and B cell identities. Mol Cell 38, 576–589, doi:10.1016/j.molcel.2010.05.004 (2010).

92 McCarthy, D. J., Chen, Y. & Smyth, G. K. Differential expression analysis of multifactor RNA-Seq experiments with respect to biological variation. Nucleic Acids Res 40, 4288–4297, doi:10.1093/nar/gks042 (2012).

93 Yang, J. et al. RBM24 is a major regulator of muscle-specific alternative splicing. Dev Cell 31, 87–99, doi:10.1016/j.devcel.2014.08.025 (2014).

94 Purcell, S. et al. PLINK: a tool set for whole-genome association and population-based linkage analyses. Am J Hum Genet 81, 559–575, doi:10.1086/519795 (2007).

