## Supplemental information for "In heart failure reactivation of RNA-binding proteins drives the transcriptome into a fetal state"

#### Supplementary Figures

#### Figure S1: Description of the subjects included in this study

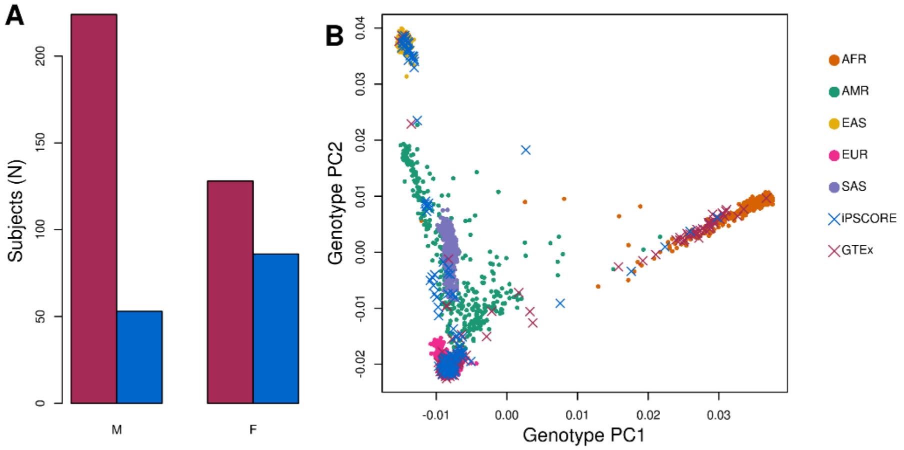


(A) Sex distribution between iPSCORE individuals (blue) and GTEx individuals (pink).

(B) PCA showing the ancestry of all subjects. Colored circles represent ancestry of individuals from the 1000 Genomes Project. GTEx and iPSCORE individuals are represented by pink and blue "X", respectively.

Details are in Table S1.

**Figure S2: Global transcriptomic differences between fetal-like iPSC-CVPC and adult CVS tissues**

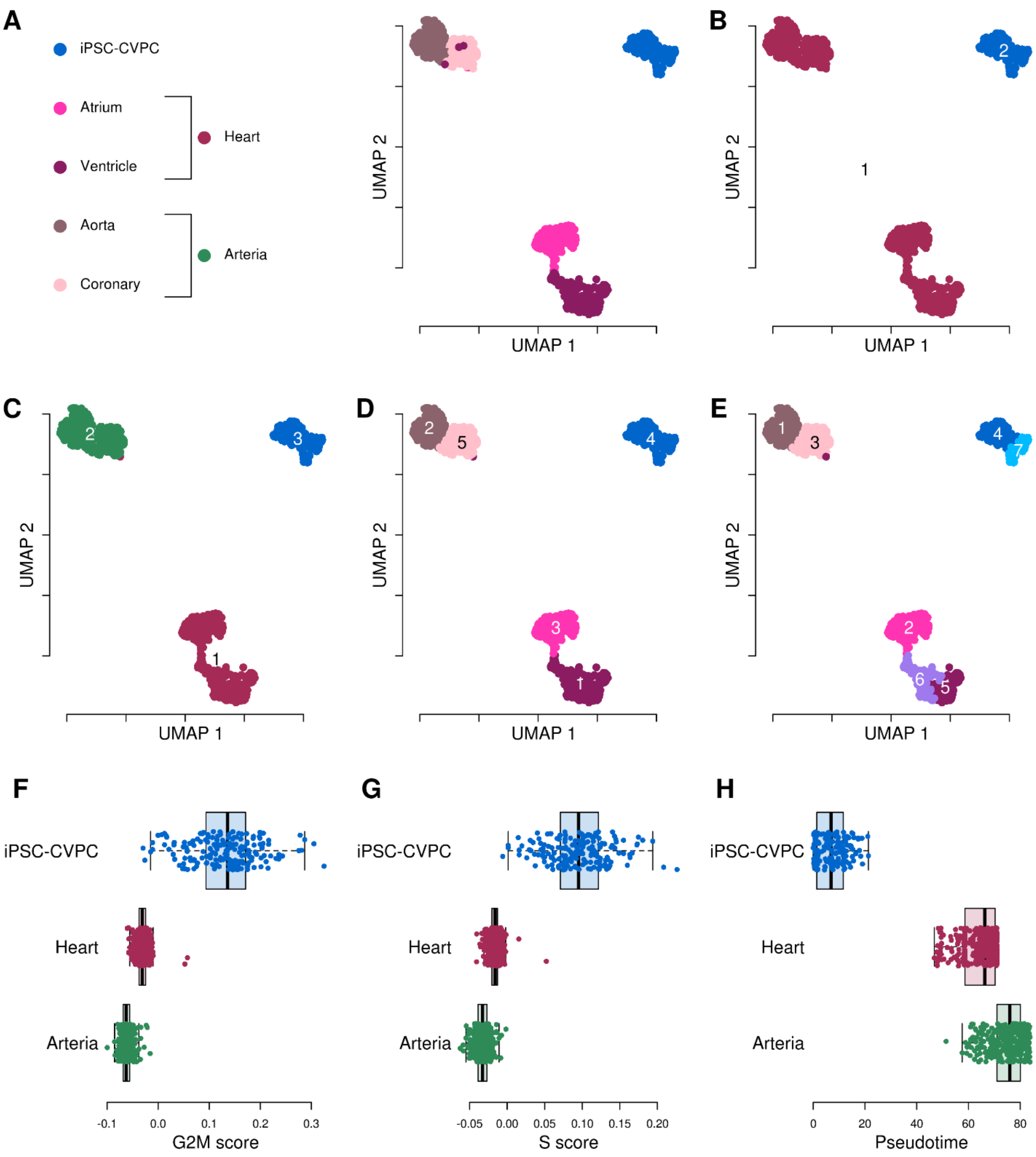


(A-E) UMAP plots showing how the 966 bulk RNA-seq CVS samples cluster at varying resolutions. Panel (A) shows the samples according to their tissue of origin (iPSC-CVPC, adult atrium, ventricle, aorta or coronary artery). In panels (B-E) samples are colored based on clustering (B) at low resolution (resolution = 0); (C) at intermediate-low (resolution = 0.01); (D) at intermediate-high resolution (resolution = 0.2); and (E) at high resolution (resolution = 0.7). Low resolution separated the samples based on their developmental stage (iPSC-CVPC and adult tissues); intermediate-low resolution divided the samples into three clusters corresponding to iPSC-CVPC, adult heart (atrial appendage and left ventricle samples) and adult arteria (aorta and coronary artery samples); intermediate-high resolution separated the samples into the five CVS tissues (iPSC-CVPC, adult atrium, ventricle, aorta or coronary artery); and high resolution separated samples derived from the same tissue-type suggesting the presence of cellular heterogeneity. For all downstream analyses, we used the clusters obtained at intermediate-low resolution. There were 5 adult heart samples that cluster with the adult arteria and 1 adult arteria sample that clusters with the adult heart. Most likely, their cell type composition differs from the other samples of the same label due to the area in which they were collected. We retained the original labeling of the 5 samples as adult heart and the 1 sample as adult arteria for downstream analyses.

(F-H) Boxplots showing (F) G2M score, (G) S phase score and (H) pseudotime calculated using Seurat or Monocle. iPSC-CVPC display a higher G2M score compare with both heart and arteria (heart: $p = 3.1 \times 10^{-83}$, arteria: $p = 1.7 \times 10^{-79}$, Mann-Whitney U test), a higher S score (heart: $p = 1.2 \times 10^{-84}$, arteria: $p = 1.6 \times 10^{-79}$, Mann-Whitney U test) and a lower pseudotime (heart: $p = 7.3 \times 10^{-85}$, arteria: $p = 1.5 \times 10^{-79}$, Mann-Whitney U test)

**Figure S3: Hierarchical clustering of 966 RNA-seq CVS samples at different resolutions**

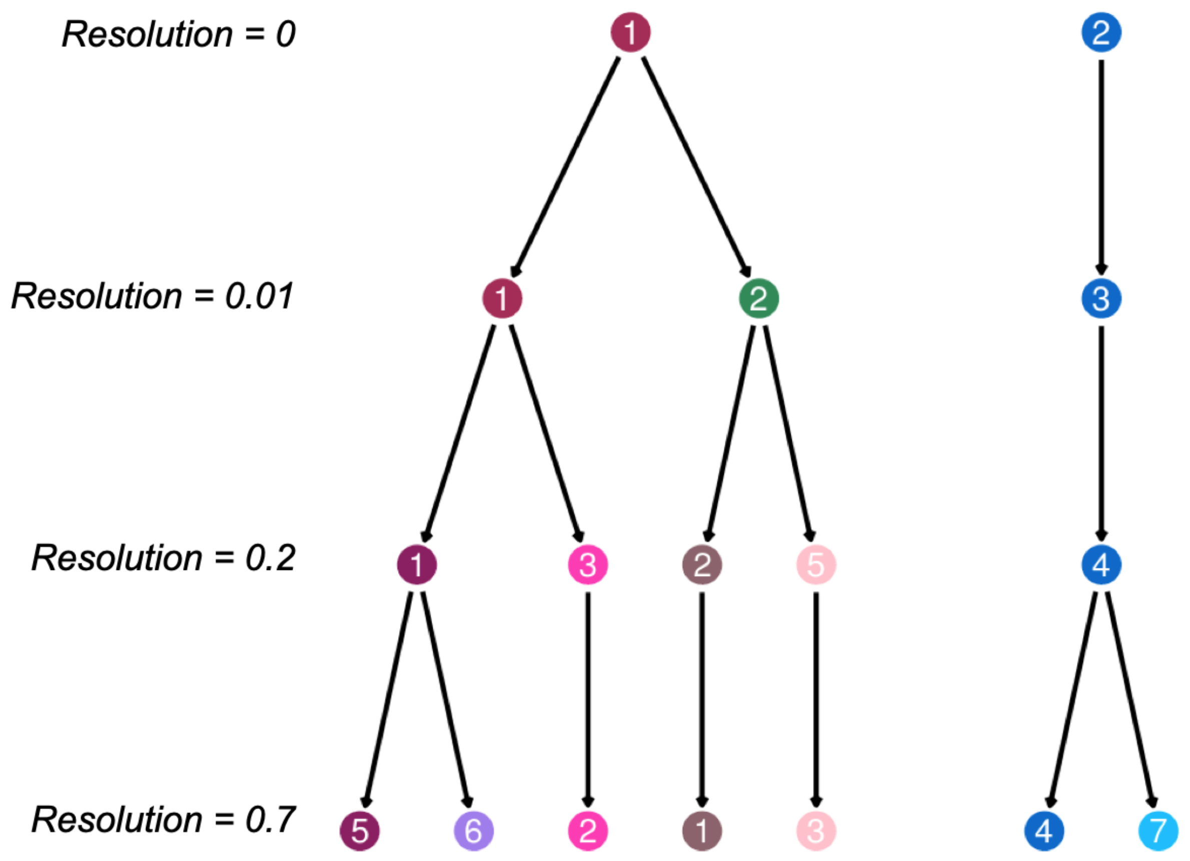


Cluster trees showing how 966 RNA-seq samples are grouped together with KNN clustering at different resolutions. The nodes are color-coded in the same manner as the clusters in the companion UMAP plots in Figure S2A-E. Lower resolution separates the samples based on developmental stage, while intermediate and higher resolutions resolve iPSC-CVPC (right) and adult CVS (left) samples into sub-clusters.

#### Figure S4: Principal components capture transcriptomic differences between CVS tissues

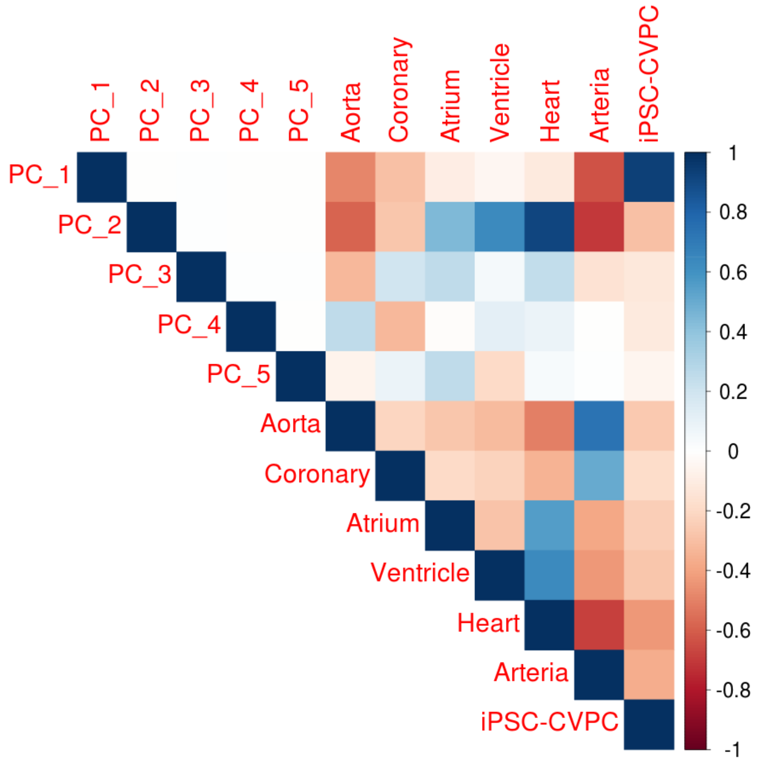


Corrplot [1] showing the Pearson correlation between the top 5 principal components calculated on the 966 CVS bulk RNA-seq samples and binary vectors indicating the type of tissue - aorta, coronary, atrium, ventricle, heart (atrium and ventricle), arteria (aorta and coronary), and iPSC-CVPC. The legend indicates the strength of the correlation for each pairwise comparison with blue representing a positive correlation and red representing a negative correlation. This correlation analysis shows that that the top principal component (PC1, 32.2% variance explained) is strongly associated with CVS developmental stage, whereas PC2 (19.4% variance explained) divides the adult CVS samples between heart and arteria. These results confirm that the two adult heart tissues (atrium and ventricle), as well as the two adult arteria tissues (aorta and coronary) have similar transcriptomes and can be combined to increase power to respectively examine "adult heart" and "adult arteria" transcriptomes. The results also show that the two adult CVS samples are more similar to each other than to the fetal-like iPSC-CVPC.

#### Figure S5: Differential expression between CVS tissues

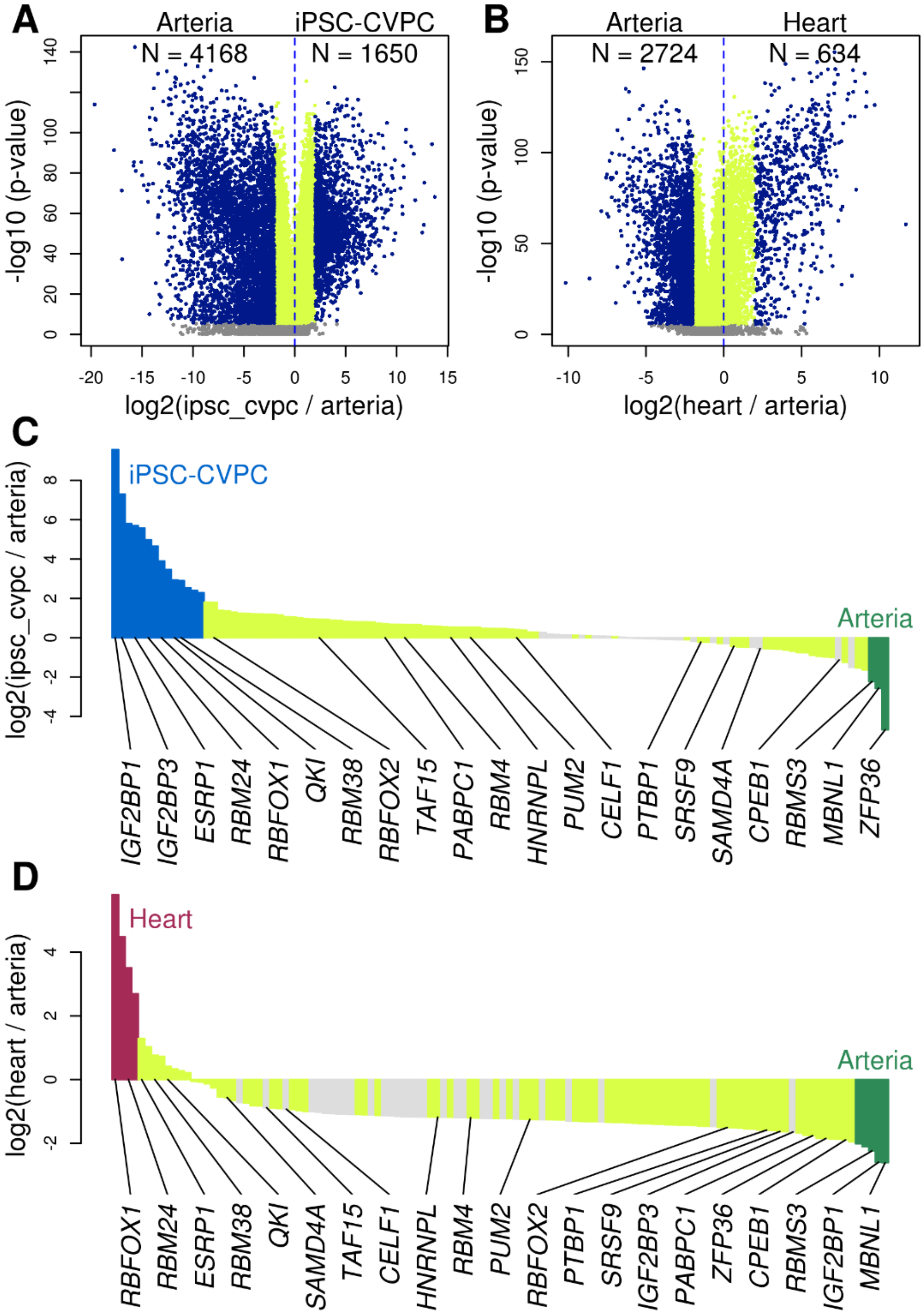


(A,B) Volcano plots showing the differentially expressed genes between (A) iPSC-CVPC and adult arteria, and (B) adult heart and adult arteria. Genes that are not differentially expressed are shown in gray, differentially expressed genes with $\log_2$ ratio between -2 and 2 are in yellow, and genes with CVS tissue-specific expression (i.e. at least four-fold expression difference; $\log_2$ ratio > 2 or < -2) are shown in blue.

(C,D) Barplots showing the $\log_2$ ratio between the mean expression in (C) iPSC-CVPC and adult arteria and (D) adult heart and adult arteria for 122 RBPs with a known motif [2-4]. RBPs shown in blue, maroon and green are iPSC-CVPC-specific, adult heart-specific and adult arteria-specific, respectively. All other differentially expressed RBPs (FDR < 0.05) are shown in yellow. These plots, in combination with Figure 1D, show that iPSC-CVPC have more overexpressed and specific RBPs than either adult heart or adult arteria, and while there is a greater number of overexpressed RBPs in adult arteria compared with adult heart, the two CVS tissues have similar numbers of tissue-specific RBPs expressed.

**Figure S6: RBPs differentially expressed between CVS tissues**

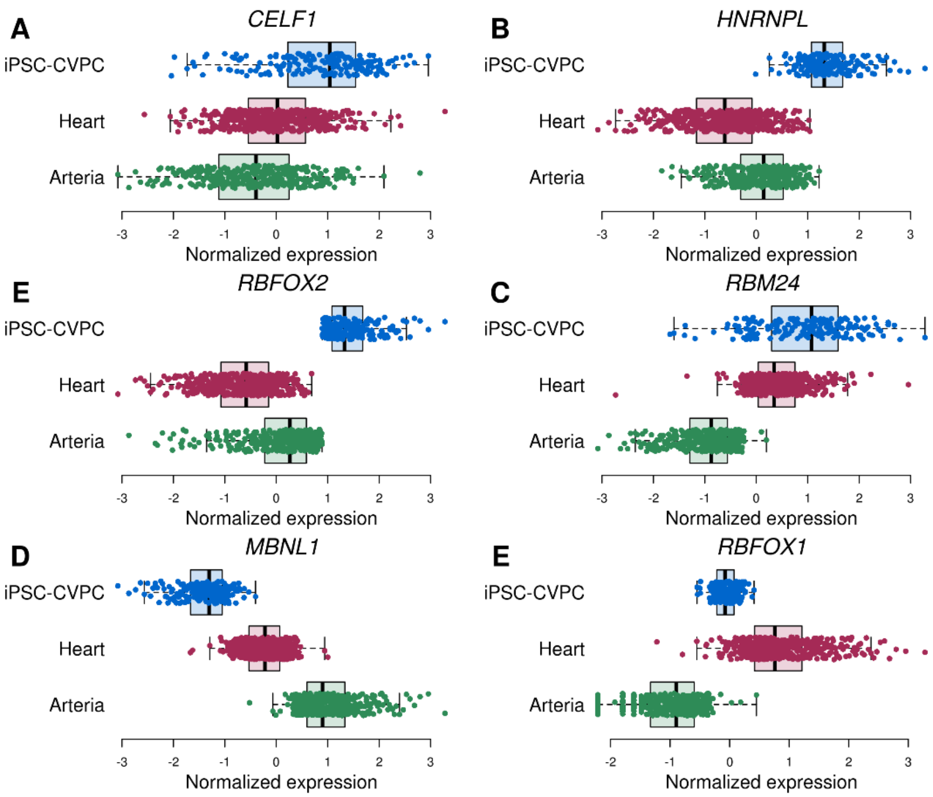


Boxplots showing gene expression levels (TPM) in iPSC-CVPC, adult heart and adult arteria for six RBPs involved in cardiac and muscle development. Four of these genes (*CELF1*, *HNRNPL*, *RBFOX2* and *RBM24*) have known functions during cardiac fetal development and were overexpressed in iPSC-CVPC compared with the two adult tissues, whereas the other two (*MBNL1* and *RBFOX1*) have adult-specific functions and were expressed at higher levels in adult arteria and adult heart, respectively, compared with iPSC-CVPC.

#### Figure S7: Differential isoform usage between iPSC-CVPC and adult heart

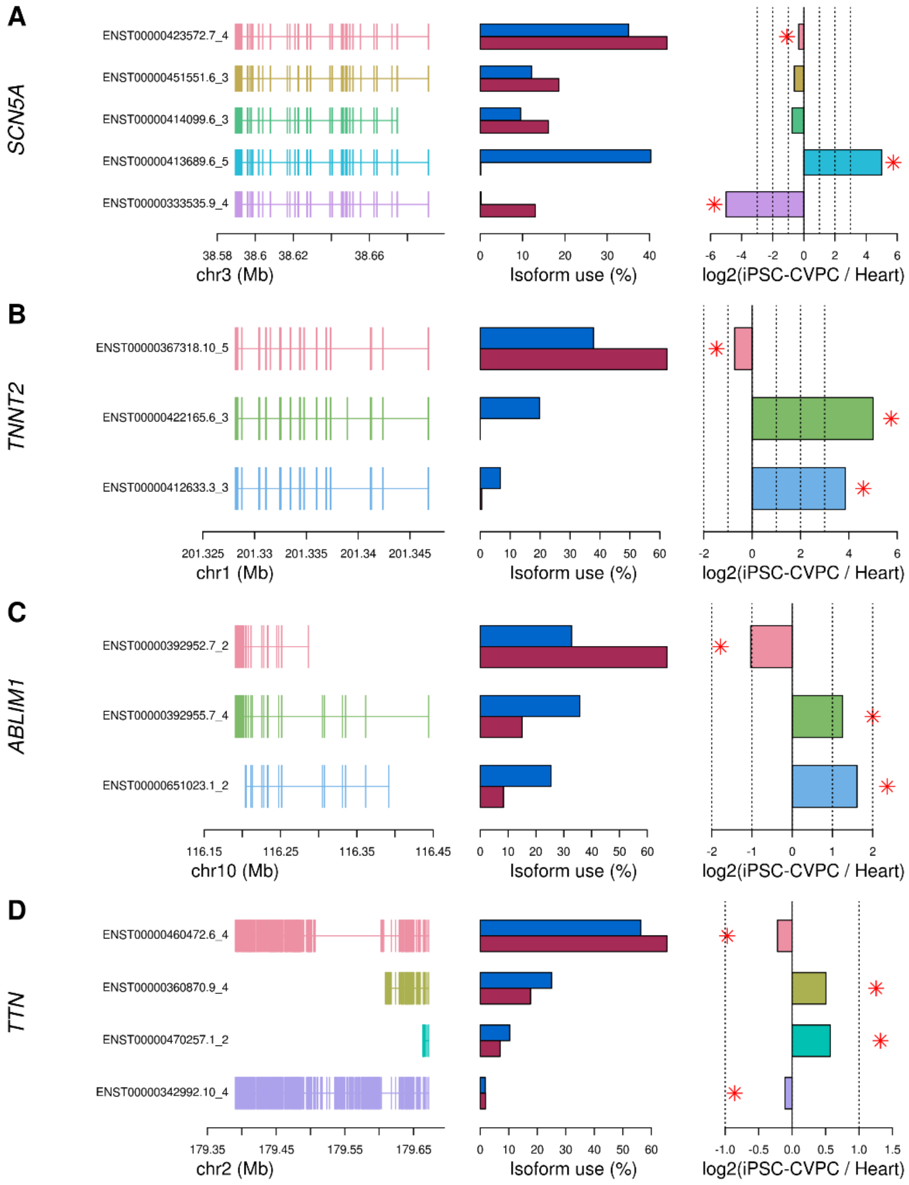


The figure shows the differential isoform usage between iPSC-CVPC (blue in the central panel) and adult heart (purple) for four cardiac genes known to have well-established developmental stage-specific isoforms: *SCN5A*, *TNNT2*, *ABLIM1* and *TTN*. Left side of the figure represents the exon structure of each expressed isoform; the barplots in the middle show the mean isoform use (%) of each isoform in iPSC-CVPC (blue) and adult heart (atrium and ventricle, purple). Barplots on the right show the $\log_2$ ratio between the mean isoform use in iPSC-CVPC and adult heart. Red stars indicate significantly differentially expressed isoforms (FDR < 0.05). Dotted lines represent one-, two- and four-fold differences between iPSC-CVPC and adult.

We found that three *SCN5A* isoforms, three *TNNT2* isoforms, three *ABLIM1* isoforms and four *TTN* isoforms were differentially expressed. Four of these isoforms (two for *SCN5A* and two for *TNNT2*) were tissue-specific, having at least a four-fold change difference between CVS tissues. To test if the isoforms expressed in the fetal-like iPSC-CVPC and in the adult heart reflect the known differences between fetal and adult, we further investigated their structure. The main differences between *SCN5A* fetal and adult isoforms are two mutually exclusive exons that encode for the voltage-sensing transmembrane domain and result in different electrophysiological properties: 6A (fetal) and 6B (adult) [5-7]. We confirmed that they were correctly expressed in the iPSC-CVPC- and adult-specific isoforms (Figure S7A). *TNNT2* isoforms differ at the 5' end, with the longer isoforms being fetal-specific and the shorter isoforms being expressed in the adult [8]. Of the three differentially expressed isoforms, we observed that the two longer isoforms were iPSC-CVPC-specific, whereas the shorter was enriched in the adult heart (Figure S7B). ABLIM1 has four C-terminal LIM domains that play a role in its binding to actin [9]. The shorter isoforms, which only contains three LIM domains because of exon 11-12 skipping, are expressed earlier during development [10]. We confirmed that the two isoforms overexpressed in iPSC-CVPC do not express these two exons (Figure S7C). Likewise, *TTN* has multiple isoforms, which accomplish different functions during embryonic development and in the adult heart: the longer fetal isoforms are responsible for the low stiffness of the fetal myocardium, whereas the short adult isoforms are associated with increased passive myocardial stiffness and may play a role in the adjustment of the cardiac muscle to increased diastolic function during development [11]. Two of the shorter *TTN* isoforms were overexpressed in the adult heart (ENST00000460472.6_4: $p = 2.6 \times 10^{-19}$, ENST00000342992.10_4: $p = 6.1 \times 10^{-33}$), whereas two of the longer isoforms were overexpressed in iPSC-CVPC (ENST00000360870.9_4: $p = 9.0 \times 10^{-17}$, ENST00000470257.1_2: $p = 2.0 \times 10^{-13}$, Figure S7D). These observations confirm that cardiac genes with established fetal-specific and adult-specific isoforms, including *SCN5A*, *TTN*, *ABLIM1* and *TNNT2* [8-18], displayed the expected differential isoform usage in the iPSC-CVPC and adult heart.

**Figure S8: Fetal-like CVS tissue has more functionally active RBPs**

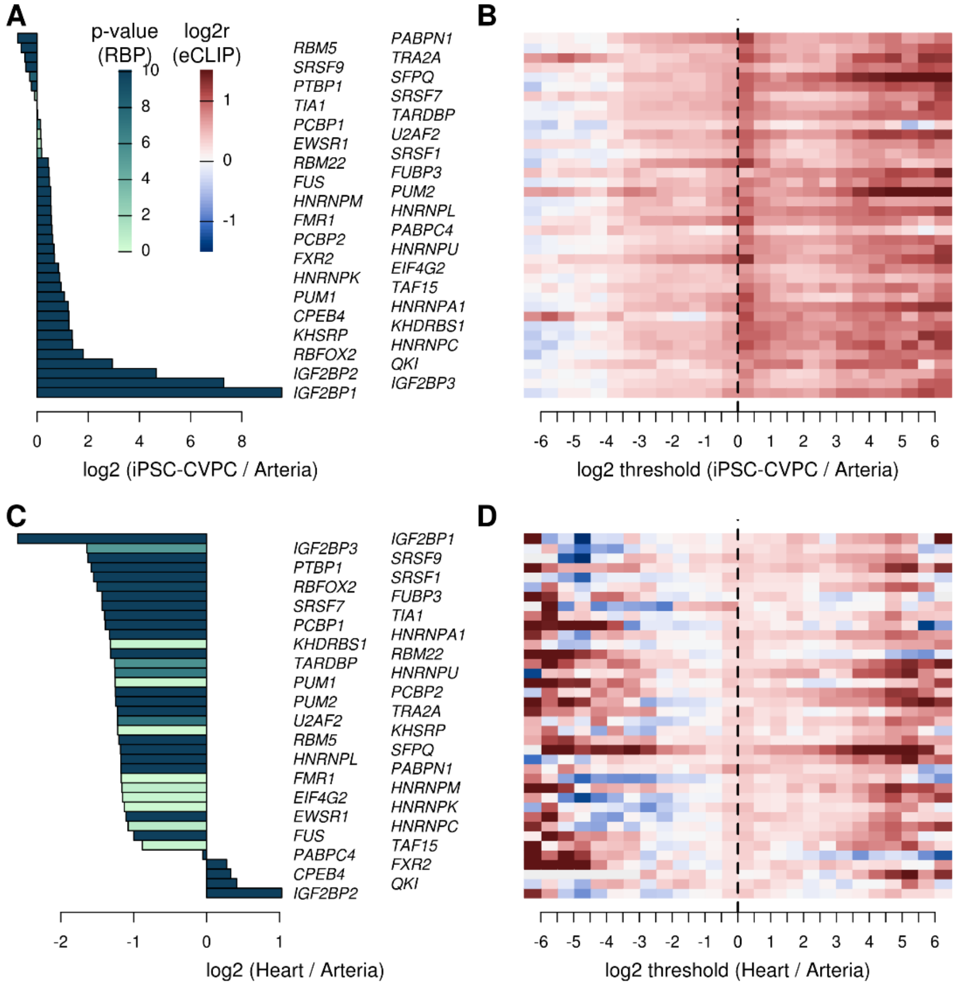


Barplots showing the differential expression of RBPs between (A) iPSC-CVPC and adult arteria and (C) adult heart and adult arteria. Colors represent $-\log_{10}$ p-values (Table S3). RBPs in panel B are sorted as in panel A, RBPs in panel D are sorted as in panel C.

(B,D) Heatmaps showing the enrichment of genomic loci encoding (B) iPSC-CVPC-specific isoforms versus adult arteria-specific isoforms and (D) adult heart-specific isoforms versus adult arteria-specific isoforms for overlapping experimentally determined (eCLIP) RBP binding sites. Each row of the heatmaps corresponds to an eCLIP experiment for the indicated RBP. Each column corresponds to $\log_2$ ratio thresholds: (B) mean isoform usage in iPSC-CVPC / mean isoform usage in adult arteria; and (D) mean isoform usage in adult heart / mean isoform usage in adult arteria. Enrichment was calculated by comparing the proportion of genes with differentially expressed isoforms passing the $\log_2$ ratio thresholds described on the X axis (Table S5) and overlapping eCLIP peaks against the proportion of genes without any differentially expressed isoforms (FDR > 0.05 for all the isoforms) overlapping eCLIP peaks. These plots show that genes with iPSC-CVPC-specific isoforms (i.e., positive thresholds in panel B) are more likely to overlap eCLIP peaks than genes with adult arteria-specific isoforms (i.e. negative thresholds in panel B); whereas adult heart-specific isoforms (i.e., positive thresholds in panel D) and adult arteria-specific isoforms (i.e. negative thresholds in panel D) are equally likely to overlap eCLIP peaks.

### Figure S9: iPSC-CVPC-specific exons enriched for splice sites overlapping RBP motifs

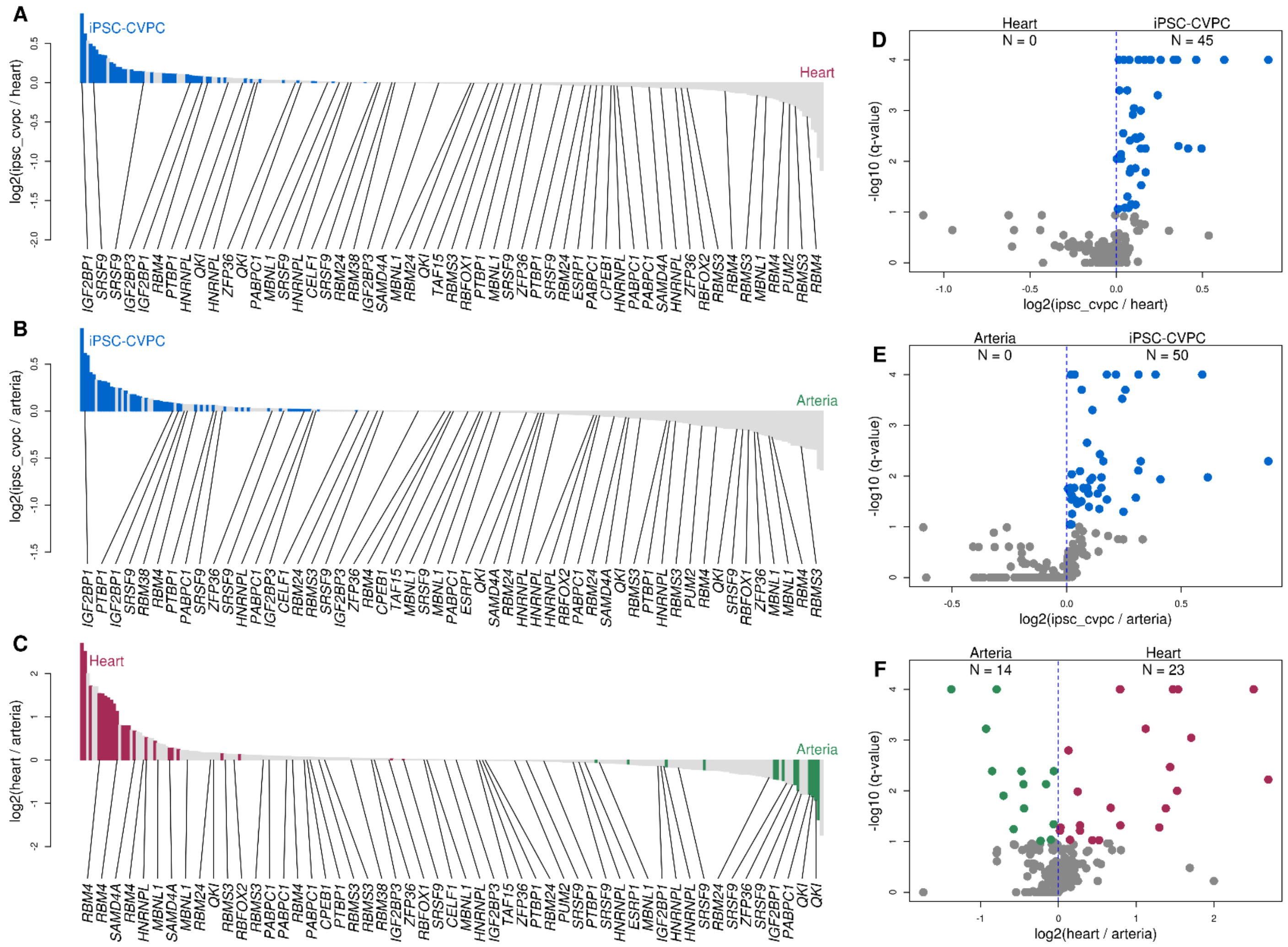


Shown are the differences in the likelihood of the 100 bp upstream of the acceptor splice sites of iPSC-CVPC-specific, adult heart-specific and adult arteria-specific exons to harbor RBP motifs.

(A-C) Barplots showing the $\log_2$ ratio of the likelihood of each tissue-specific splice site to harbor a RBP motif (iPSC-CVPC = blue, adult heart = red, adult arteria = green).

(D-F) Volcano plots show the enrichment for each RBP (X axis, same as panels A-C) and its associated $-\log_{10}$(q-value) calculated using Homer *findMotifsGenome.pl* [19]. Q-values < $1.0 \times 10^{-5}$ are shown as $1.0 \times 10^{-5}$. Colors are as shown in panels A-C.

**Figure S10: Clustering of FACs-sorted cardiac cells from *Tabula Muris***

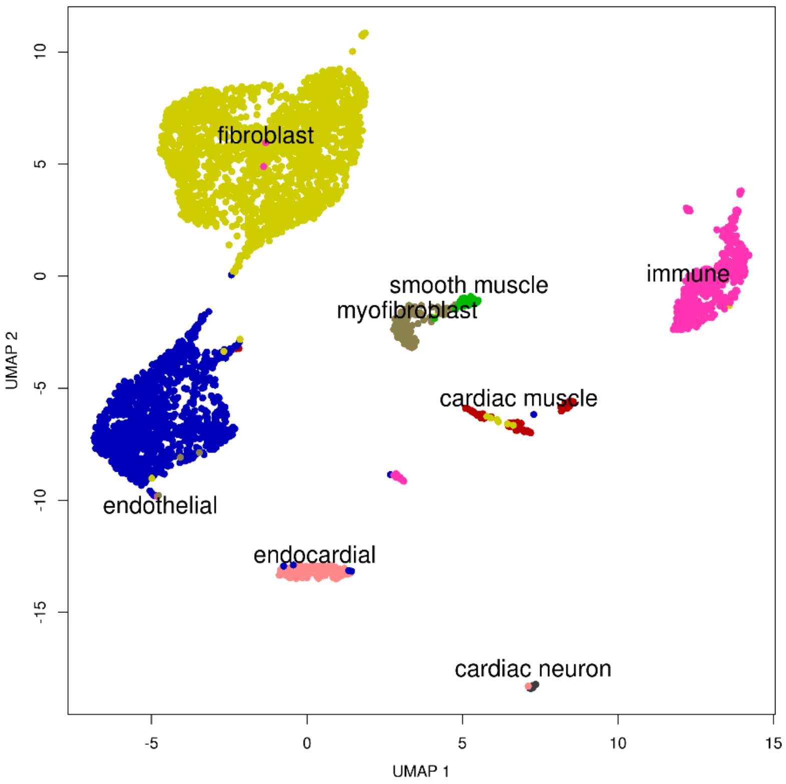


UMAP plot showing the clustering and cell types associated with each of the 4,365 cells obtained from *Tabula Muris*. Cell labels are as defined by the *Tabula Muris* Consortium [20].

#### Figure S11: Principal components capture cellular heterogeneity across 966 CVS samples

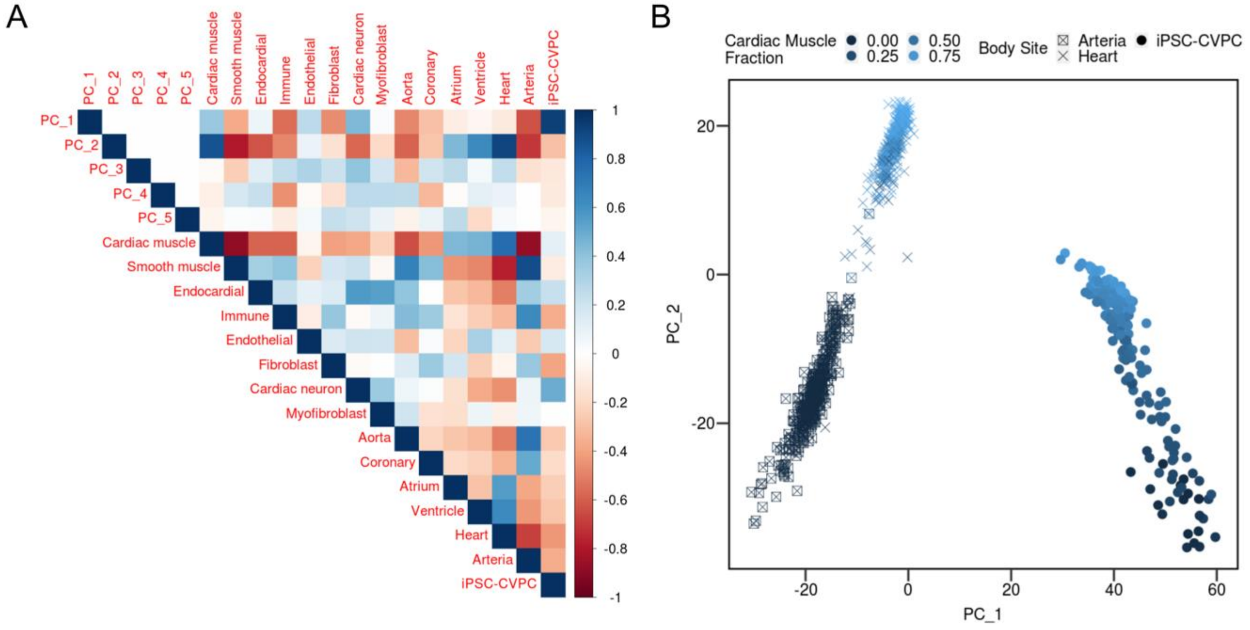


A) Corrplot showing the Pearson correlations between the top 5 principal components calculated on 2,000 most variable genes across the 966 bulk RNA-seq CVS samples (Figure 1B-F), cell type proportion, and the tissue types of the 966 CVS samples. The top two principal components were correlated with the cardiac muscle proportion (R = 0.3749 and 0.8643, respectively), indicating that gene expression levels are driven by cellular heterogeneity and suggesting that the heterogeneity observed at high clustering resolution (Figure S2E) is a consequence of the variability in cell type proportions within each tissue.

B) Scatter plot showing the separation of fetal iPSC-CVPC (right) and adult heart and adult arteria (left) by the first principal component and the separation of samples with larger fraction of cardiac muscle (top / light blue) and those with smaller fraction (bottom / dark blue) by the second principal component. These results further highlight that developmental stage and cellular heterogeneity explain the most variability in the transcriptomes of the 966 CVS samples.

#### Figure S12: Associations between gene expression and cell type proportions

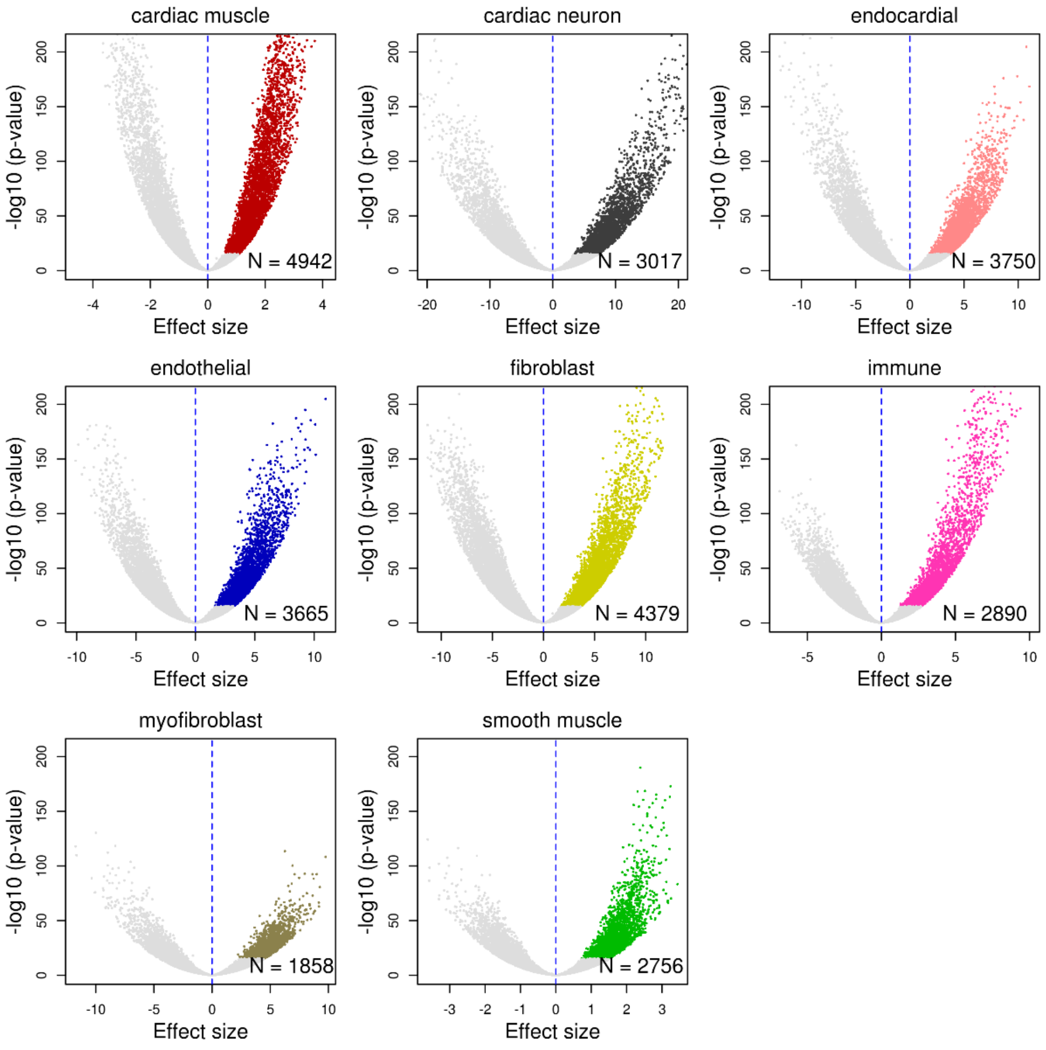


Volcano plots showing the number of genes whose expression is associated with the proportion of indicated cell type. All non-significant genes are colored in light gray (FDR > 0.05). Genes significantly associated with each cell type (FDR < 0.05 and effect size > 0) are colored and their number is reported at the bottom-right of each plot.

#### Figure S13: Validating cell type associated gene expression with scRNA-seq

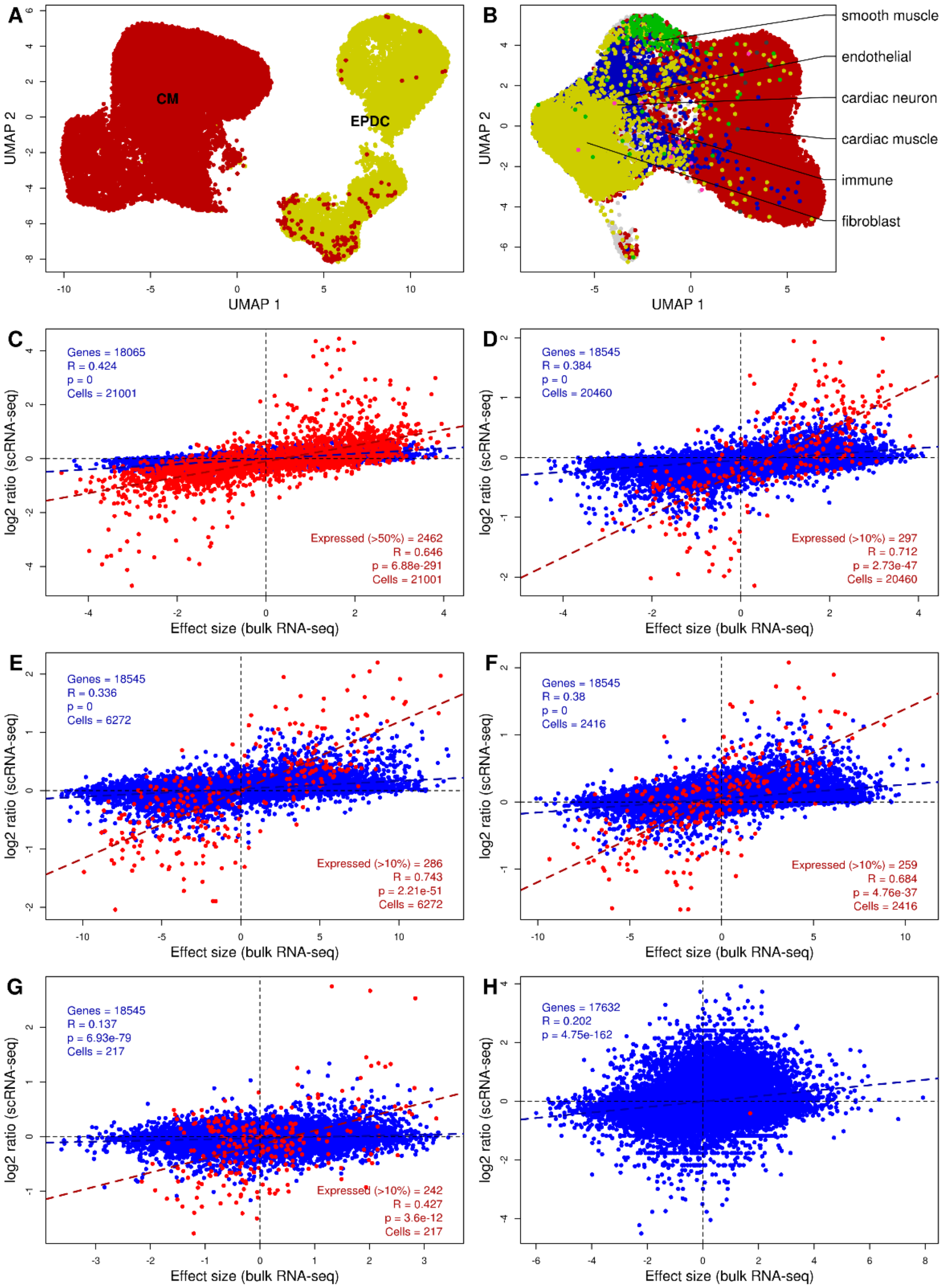


We used two recent scRNA-seq datasets from iPSC-CVPC and adult heart [21,22] to further characterize the cell type proportions estimated by CIBERSORT. Using scRNA-seq data, we performed differential expression analysis for each cell type and found that the ratio between the expression levels of each gene in a cell type compared with all the other cells was strongly correlated with the effect size of the associations between estimated cell type proportions determined by CIBERSORT deconvolution and gene expression across all cell types.

(A-B) Plots showing the UMAP coordinates of (A) 32,026 iPSC-CVPC cells from eight iPSCORE samples; and (B) 33,050 adult left ventricle cells.

(C) Scatterplot showing the effect size of the association between gene expression and % of cardiac muscle in the 966 deconvoluted RNA-seq CVS samples (X axis) and the $\log_2$ ratio between the expression in cardiomyocytes (CMs) and EPDCs in the 32,026 iPSC-CVPC cells (Y axis).

(D-G) Scatterplots showing the effect size of the association between gene expression and % of (D) cardiac muscle, (E) fibroblast, (F) endothelial and (G) smooth muscle in the 966 deconvoluted RNA-seq CVS samples (X axis) and the $\log_2$ ratio between gene expression in the indicated cell type and all other cells for the 33,050 adult left ventricle cells. Red shows highly expressed genes (expressed in at least 50% of iPSC-CVPC cells or 10% of adult cells). All the lowly expressed genes are shown in blue. Red and blue dashed lines are the regression lines for highly expressed genes and across all tested genes, respectively. In the bottom right corner of each plot, the correlation for the highly expressed genes is shown. In the top left corner, the correlation calculated across all the genes is shown. These plots show that the associations between gene expression and the estimated cell type proportion in the bulk RNA-seq data are significantly strongly correlated with the expression differences observed in the scRNA-seq data, both in iPSC-CVPC and adult heart, indicating that cellular deconvolution and differential gene expression analysis reflect differences in cell composition between samples.

(H) Scatterplot showing the difference between effect sizes of the association between cardiac muscle proportion and gene expression between iPSC-CVPC and adult heart (X axis) and the $\log_2$ ratio between average normalized expression in iPSC-CVPC and adult cardiac muscle cells. The strong positive correlation indicates that the stage-specific associations in cardiac muscle between iPSC-CVPC and adult heart reflect gene expression differences between iPSC-CVPC and adult cardiac muscle cells observed using scRNA-seq.

#### Figure S14: Differential isoform expression: linear model versus ridge regression

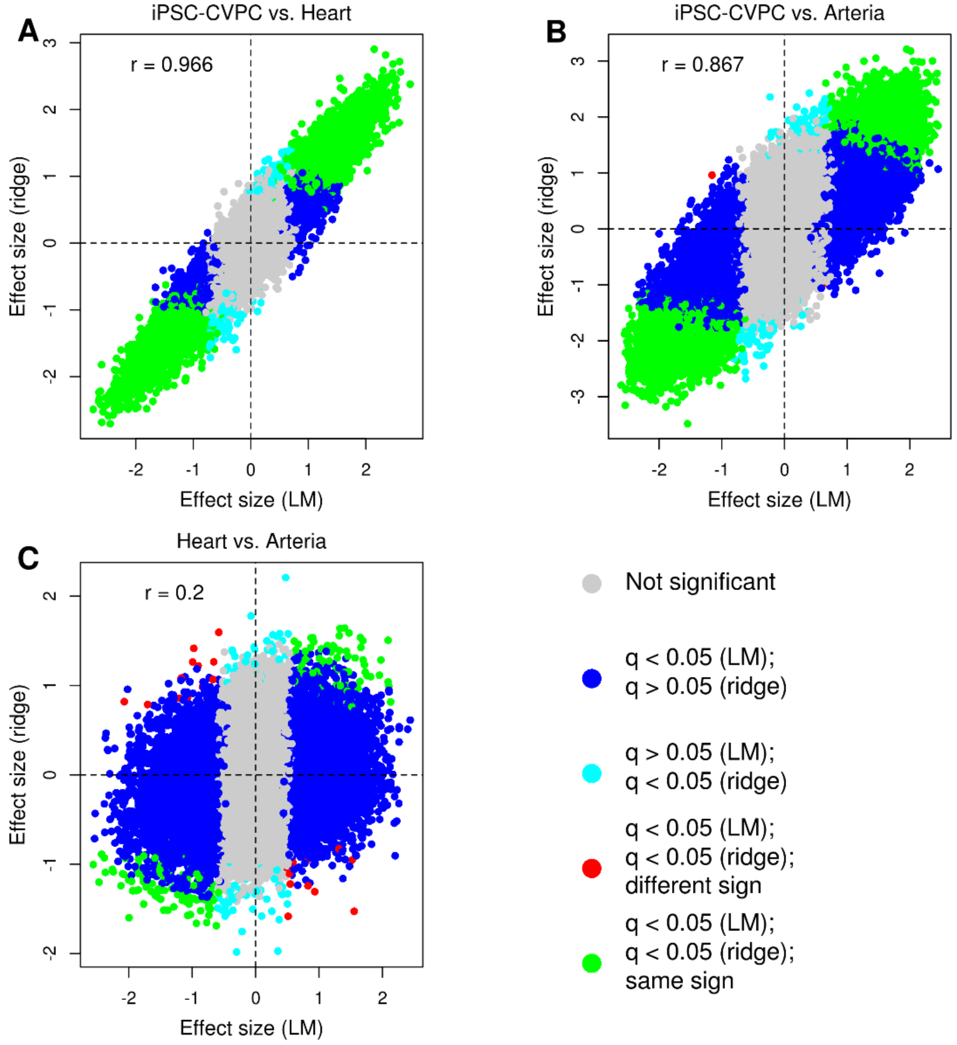


Scatterplots showing the differential isoform usage effect size without considering cell types (linear model, LM, X axis) and considering the cell type proportions as covariates (ridge regression, Y axis) for each pairwise comparison between CVS tissues: (A) iPSC-CVPC vs. adult heart; (B) iPSC-CVPC vs. adult arteria; and (C) adult heart vs. adult arteria. Isoforms differentially expressed in both analyses and that have the same effect size sign are shown in green; isoforms whose effect size has different signs between the LM and ridge regression are in red; isoforms that are differentially expressed only using the LM are shown in blue; and isoforms that are differentially expressed only when taking cell type proportions into account (ridge) are in cyan.

#### Figure S15: Gene expression associations with both CVS tissues and cell type proportions

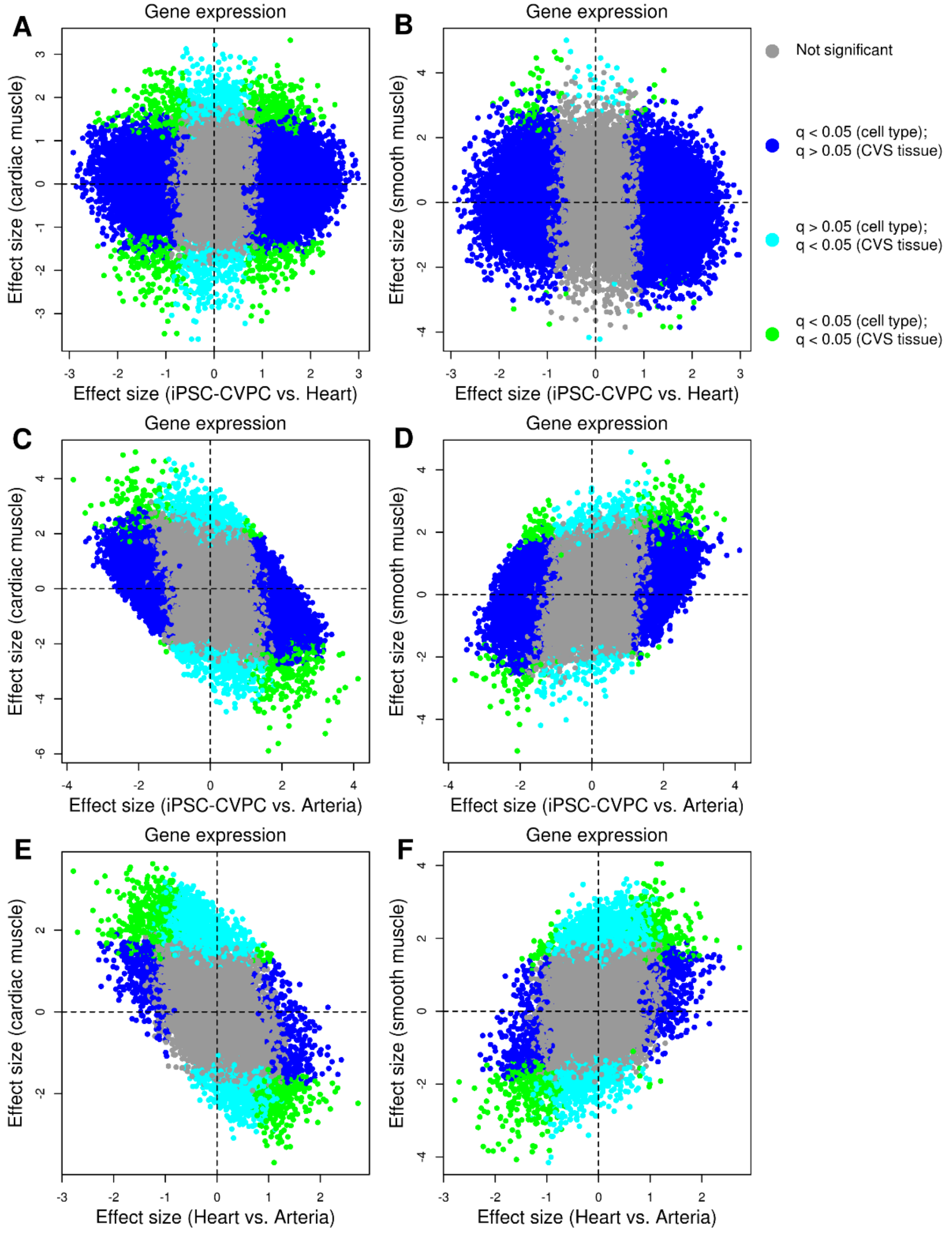


Scatterplots showing differential gene expression effect size between each indicated pair of CVS tissues and the effect size of the association between gene expression and cell type proportion of cardiac muscle or smooth muscle cells. Genes whose expression are not associated with either CVS tissue or cell type proportion are shown in gray; genes significantly associated with both CVS tissue and cell type proportion are in green; genes associated only with cell type are in blue; and genes that are associated only with CVS tissue are in cyan.

#### Figure S16: RBPs associated with CVS tissues and cell type proportions

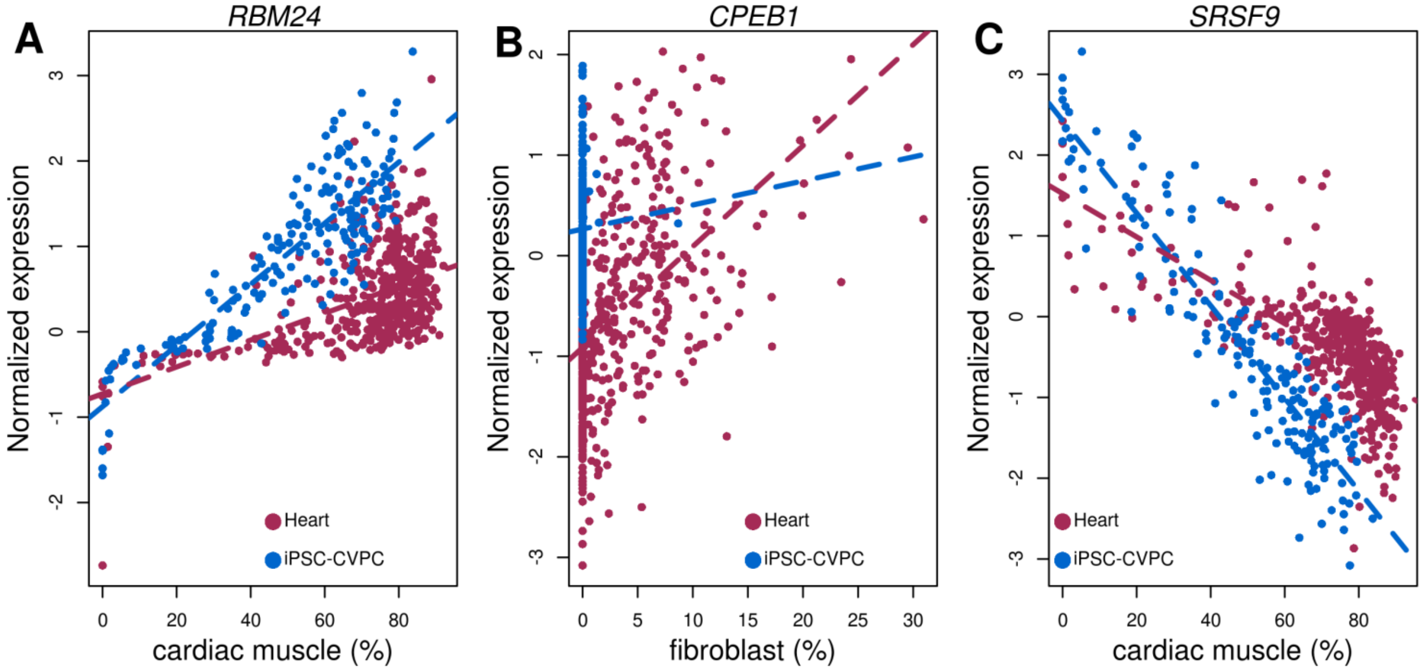


Scatterplots showing cell type proportions (X axis) and normalized expression in iPSC-CVPC and adult heart (Y axis) for the RBPs that were both differentially expressed between iPSC-CVPC and adult heart (FDR < 0.05) and positively associated with at least one cell type (effect size > 0 and FDR < 0.05). Dashed lines represent regression lines calculated on each CVS tissue.

*RBM24*, a major regulator of muscle-specific alternative splicing [23], regulates apoptosis during development and its deficiency is associated with embryonic lethality due to major cardiac malformations in mice [24]. It was significantly overexpressed in iPSC-CVPC ($p = 6.7 \times 10^{-27}$, ridge regression) and positively associated with the cardiac muscle proportion ($p = 3.8 \times 10^{-7}$, Figure S16A, Table S15). Similarly, *CPEB1*, which is involved in germ cell differentiation and regulates senescence in fibroblasts [25,26], was overexpressed in iPSC-CVPC ($p = 2.3 \times 10^{-7}$) and positively associated with the fibroblast proportion ($p = 1.7 \times 10^{-37}$, Figure S16B). Conversely, *SRSF9*, whose expression impairs muscle differentiation [27], was overexpressed in the adult heart ($p = 1.1 \times 10^{-33}$) and negatively associated with the cardiac muscle proportion ($p = 2.9 \times 10^{-35}$, Figure S16C).

### Supplementary Tables

#### Table S1: Description of subjects included in this study

The table show subject information for all the 491 individuals included in this study, including: subject ID; subject name; source (iPSCORE or GTEx); sex; age, height; weight; BMI; and 20 genotype principal components. Columns AC-AG represent the family information for each iPSCORE subject, as included in dbGaP (phs001325) as part of the iPSCORE Resource: “family ID” classifies the subject by family to identify related family members; “twin ID” identifies the dbGaP ID if the subject is a twin; “twin type” indicates the type of twin (MZ = monozygotic; DZ = dizygotic) if the subject is a twin; “father ID” and “mother ID” indicate the subject_UUID of the father and mother of the subject if part of the iPSCORE resource.

#### Table S2: Description of bulk RNA-seq samples

Shown are the 966 bulk RNA-seq samples used in this study, including: source (iPSCORE or GTEx); subject ID; assay ID; SRA run ID for the GTEx RNA-seq samples downloaded from dbGaP; iPSCORE unique differentiation identified (UDID); iPSCORE iPSC line identifier submitted to dbGap (phs001325), which indicates clone and passage of iPSC; iPSCORE iPSC ID; total number of reads; % uniquely mapped reads; % of mitochondrial reads, calculated as the number of reads mapping to mitochondrial genes divided by the total number of reads mapping to genes; tissue and organ associated with each sample (arteria, heart, iPSC-CVPC, aorta, coronary artery, atrial appendage or left ventricle: 0 = absent; 1 = present); ten principal components calculated on the expression of 2,000 genes across the 966 CVS samples using Seurat; UMAP coordinates of each sample after clustering by Seurat; G2M and S phase score calculated using Seurat; estimated pseudotime of each sample using Monocle; cluster membership of each sample calculated using Seurat at four different resolutions (Figure 1A-B, Figure S2, Figure S3); estimated cell type proportions deconvoluted using CIBERSORT; the last column (“Trimmed”) indicates which iPSCORE samples had their read length trimmed to 75 bp to test whether different read lengths between iPSCORE and GTEx affect differential gene and isoform expression analyses.

#### Table S3: Differential gene expression analysis

Shown is the differential expression analysis between: iPSC-CVPC and adult heart; iPSC-CVPC and adult arteria; adult heart and adult arteria. For each gene, we report: gene ID; gene name; the tested tissues (tissue 1 and tissue 2); effect size, standard error, p-value and FDR correction (Bonferroni). Effect size > 0 corresponds to genes where the expression in tissue 1 is greater than tissue 2. Only differentially expressed genes are reported (FDR < 0.05). The differential expression analysis of all genes can be found at https://figshare.com/s/b2e70a2ba3a4aac935d9.

#### Table S4: Functional enrichment analysis (genes)

The table shows the functional enrichment analysis for genes differentially expressed between each pair of CVS tissues (Table S3). For each gene set, we report: the tested tissues (tissue 1 and tissue 2); the gene set collection, as defined by MSigDB, the gene set name and its URL; the number of tested genes in the gene set; the average effect size for all the tested genes in the gene set and for all the other expressed genes; p-value (t-test); and FDR (Benjamini-Hochberg). Only significant gene sets are reported (FDR < 0.05). The differential expression analysis of gene sets can be found at https://figshare.com/s/b2e70a2ba3a4aac935d9.

#### Table S5: Differential isoform expression analysis

Shown is the differential isoform expression analysis between: iPSC-CVPC and adult heart; iPSC-CVPC and adult arteria; adult heart and adult arteria. For each isoform, we report: isoform ID, gene ID, gene name; the tested tissues (tissue 1 and tissue 2); effect size, standard error, p-value and FDR correction (Bonferroni); mean isoform use in each of the two tested tissues and the $\log_2$ ratio between them; whether the isoform is differentially expressed (FDR < 0.05). Effect size > 0 corresponds to isoforms where the expression in tissue 1 is greater than tissue 2. Only CVS tissue-specific isoforms are reported (FDR < 0.05 and absolute value of $\log_2$ ratio >2). The differential expression analysis of all isoforms can be found at https://figshare.com/s/b2e70a2ba3a4aac935d9.

#### Table S6: Functional enrichment analysis (isoforms)

The table shows the functional enrichment analysis of genes that have at least one CVS tissue-specific isoform, compared with genes that do not have differentially expressed isoforms between each pair of CVS tissues (Table S5). For each gene set, we report: the tested tissues (tissue 1 and tissue 2); the gene set collection, as defined by MSigDB, the gene set name and its URL; the number of tested genes in the gene set; the enrichment calculated using the *estimate* parameter in the *fisher.test* function in R; p-value (Fisher's exact test); and FDR (Benjamini-Hochberg). Only significant gene sets are reported (FDR < 0.05). The differential expression analysis of gene sets can be found at https://figshare.com/s/b2e70a2ba3a4aac935d9.

#### Table S7: Overlap of each CVS tissue-specific exon with protein domains

The table shows the overlap between each CVS tissue-specific exon and protein domains obtained from the Prot2HG database [28]. For each CVS tissue-specific exon, we report: the exon ID, transcript ID, gene ID, gene name; the tested tissues ("iPSC-CVPC vs. heart", "iPSC-CVPC vs. arteria" or "heart vs. arteria"); the tested CVS tissue the exon is expressed in, and the overlapping protein domain of the exon.

#### Table S8: Enrichment of CVS tissue-specific isoforms for RBP binding sites

For each pairwise comparison between CVS tissues (columns A and B), the table shows the differences in the overlap with each eCLIP experiment [3] between the gene bodies of isoforms specific to one of the two tested CVS tissues and the gene bodies of isoforms that are not differentially expressed. Enrichments were performed using increasingly stringent thresholds for determining CVS tissue-specific isoforms, based on the $\log_2$ ratio between the mean isoform usage in the two tested CVS tissues. The table shows: 1) the two tested CVS tissues (column A); the tested RBP, its associated eCLIP experiment and the cell line in which the eCLIP experiment was performed (columns B-D); the $\log_2$ ratio threshold, as described in Table S5 (column E); and the results obtained from Fisher's exact test (*fisher.test* function in R), including the estimate (odds ratio), the $\log_2$ of the estimate, and the p-value. Values displayed in this table were used to build Figure 3B, 2H-I and Figure S8.

#### Table S9: Enrichment of CVS tissue-specific exons for RBP motifs

Tables showing the RBP motif enrichment results obtained using Homer *findMotifsGenome.pl* with CVS tissue-specific exons and 262 RBP motifs. The table shows: RBP name and motif ID (columns A,B); the tested tissues (columns C,D); $\log_2$ ratio between the two tissues calculated by Homer (column E); p-value and q-value calculated by Homer (columns F,G).

#### Table S10: Enrichment of CVS tissue-specific exons for canonical splice sites

For each pairwise comparison between CVS tissues, (column A,B: iPSC-CVPC vs. adult heart, iPSC-CVPC vs. adult arteria and adult heart vs. adult arteria), the table shows the differences in the probability of each base pair occurring in the 100 bp upstream of the splice acceptor site and the 100 bp downstream of the splice donor site. The table shows: the two tested CVS tissues (columns A, B); which splice site is tested (donor or acceptor, column C); the tested nucleotide (column D); the position relative to the exon start (negative values, upstream of the splice acceptor site) or end (positive values, downstream of the splice donor site) (column E); and the results obtained from Fisher's exact test (*fisher.test* function in R), including the estimate (odds ratio, column F), its 95% confidence interval (column G,H) and the p-value (column I). In Figure 3C-I, only the tests associated with the most common nucleotide at each position are shown.

#### Table S11: Marker genes for cardiac cell types in *Tabula Muris*

Shown are marker genes identified for each cluster using the *FindAllMarkers* function in Seurat. Column C represents the fraction of cells where the gene described in column A is expressed in the cell type described in column B; column C represents the % of cells in the cells labeled as the cell type in column B that express the gene; column D represents the % of cells in all other cell types that express the same gene; column E shows the $\log_2$ fold change between the average expression level in the cell type described in column B and all other cells; column F and G represent the p-value (Wilcoxon rank sum test) and FDR-adjusted p-value (Bonferroni). All values described in this table were calculated using the *FindAllMarkers* function in Seurat with parameters *min.pct = 0.2*, *logfc.threshold = 0.1*. Only genes with FDR < 0.1 are shown. If a gene is significantly overexpressed in more than one cell type, it is reported multiple times.

#### Table S12: Expression matrix input to CIBERSORT

Shown is the average expression of all marker genes described in Table S11. Expression levels were obtained using the Seurat function *AverageExpression*. This table was used as input for CIBERSORT. CIBERSORT results are reported in Table S2 with all the variables used as covariates for the differential expression analysis.

#### Table S13: Differential gene expression analysis (cell types)

Shown are the associations between the expression of each expressed gene and isoform and cell type proportions in all bulk RNA-seq CVS samples. For each gene and isoform, we report: isoform ID, gene ID, gene name; the cell type whose proportion is tested for association with expression; effect size, standard error, p-value, FDR correction (Bonferroni); whether the expression of the gene or isoform is associated with cell type proportion (FDR < 0.05 and effect size > 0). Only differentially expressed genes and isoforms are reported (FDR < 0.05). The differential expression analysis of all genes and isoforms can be found at https://figshare.com/s/b2e70a2ba3a4aac935d9.

#### Table S14: Functional enrichment analysis (cell types)

The table shows the functional enrichment analysis for genes associated with each cell type. The table is organized as Table S4, with a difference: cell type, rather than "tissue 1" and "tissue 2" is reported. For each cell type, we determined the association (effect size) between its proportion and the expression of each gene, and used the effect sizes of all expressed genes as input for gene set enrichment analysis (GSEA). We observed that the most significantly enriched gene sets corresponded to the main function associated with each cell type, including mitochondrial and cell respiration functions for cardiac muscle, immune response for immune cells, cytoskeleton and actin binding for smooth muscle. Only significant gene sets are reported (FDR < 0.05). The differential expression analysis of gene sets can be found at https://figshare.com/s/b2e70a2ba3a4aac935d9.

#### Table S15: Differential gene and isoform expression using cell type proportions as covariates

Shown is the differential expression analysis for both genes and isoforms performed using ridge regression and cell type proportions as covariates between: iPSC-CVPC and adult heart; iPSC-CVPC and adult arteria; adult heart and adult arteria. For each gene and isoform, we report: isoform ID, gene ID and gene name; the tested tissues (tissue 1 and tissue 2); the covariate, including "tissue", which represents the differential expression between tissue 1 and tissue 2, and each of the cell types; effect size, standard error, p-value and FDR correction (Bonferroni). Only significant associations (FDR < 0.05) are reported. All tests can be found at https://figshare.com/s/b2e70a2ba3a4aac935d9.

#### Table S16: PCA of heart failure, iPSC-CVPC, adult heart and adult arteria bulk RNA-seq samples

The table shows sample information for each of the 30-heart failure bulk RNA-seq samples (15 pairs of matched pre- and post-LVAD samples from GSE46224) and the principal components coordinates of these samples, as well as the 966 iPSCORE and GTEx CVS samples. Columns A-C show the sample ID, tissue and study for all the 996 analyzed samples.

Columns D-F show subject ID, age, sex and LVAD status (pre- or post-LVAD) for each of the 30 heart failure samples. The subsequent columns show the PCA coordinates of all the samples based on gene expression.

#### Table S17: Cell proportion differences between heart failure and healthy CVS tissues

The table shows the cell proportion differences between the two heart failure sample sets (pre- and post-LVAD) and the three healthy CVS tissues (iPSC-CVPC, adult heart and adult arteria). For each pair of tissues (columns A and B) and cell type (column C), the mean cell type proportion in each of the two tissues is shown (columns D and E), as well as the p-value (t-test, column F) and its FDR correction (Benjamini-Hochberg's method, column G). We found that pre-LVAD samples had a significantly higher proportion of myofibroblasts than post-LVAD samples (2.1%, compared with 1.2%, $p = 0.0010$, t-test). All the other cell type proportion differences between pre- and post-LVAD cell type proportions were not significant after FDR correction.

#### Table S18: Differential expression between heart failure and healthy CVS tissues

The table shows the results from differential expression analysis of genes and isoforms between pre-LVAD samples and the three healthy CVS tissues (iPSC-CVPC, adult heart and adult arteria). Columns A-D show gene and transcript information, including: transcript ID, gene ID, gene name, and analysis type (gene or isoform). Columns E-F show the pairs of tested tissues. Columns G-J show the effect size, its standard error, p-value and FDR correction (Bonferroni's method). FDR correction was performed independently on genes and isoforms. All genes and isoforms with $FDR < 0.05$ were considered as significant and shown in this table. All tests can be found at https://figshare.com/s/b2e70a2ba3a4aac935d9.

#### Table S19: Genes and isoforms whose expression is associated with read length

The table shows the gene expression and isoform use differences between GTEx RNA-seq samples with short (75 bp) and iPSCORE RNA-seq samples with long (150 bp) read length. For each gene or isoform, shown are the gene or isoform ID, p-value (paired t-test), Bonferroni-corrected p-value, source (gene or isoform), gene ID if the source is isoform, and whether the gene or isoform is blacklisted.

### References


1 Wei, T. & Simko, V. R package "corrplot": Visualization of a Correlation Matrix (Version 0.84). *CRAN* (2017).

2 Paz, I., Kosti, I., Ares, M., Jr., Cline, M. & Mandel-Gutfreund, Y. RBPmap: a web server for mapping binding sites of RNA-binding proteins. *Nucleic Acids Res* **42**, W361-367, doi:10.1093/nar/gku406 (2014).

3 Van Nostrand, E. L. *et al.* A large-scale binding and functional map of human RNA-binding proteins. *Nature* **583**, 711-719, doi:10.1038/s41586-020-2077-3 (2020).

4 Ray, D. *et al.* A compendium of RNA-binding motifs for decoding gene regulation. *Nature* **499**, 172-177, doi:10.1038/nature12311 (2013).

5 Wahbi, K. *et al.* Brugada syndrome and abnormal splicing of SCN5A in myotonic dystrophy type 1. *Arch Cardiovasc Dis* **106**, 635-643, doi:10.1016/j.acvd.2013.08.003 (2013).

6 Shen, C. *et al.* Novel idiopathic DCM-related SCN5A variants localised in DI-S4 predispose electrical disorders by reducing peak sodium current density. *J Med Genet* **54**, 762-770, doi:10.1136/jmedgenet-2017-104780 (2017).

7 Freyermuth, F. *et al.* Splicing misregulation of SCN5A contributes to cardiac-conduction delay and heart arrhythmia in myotonic dystrophy. *Nat Commun* **7**, 11067, doi:10.1038/ncomms11067 (2016).

8 Gomes, A. V. *et al.* Cardiac troponin T isoforms affect the Ca(2+) sensitivity of force development in the presence of slow skeletal troponin I: insights into the role of troponin T isoforms in the fetal heart. *J Biol Chem* **279**, 49579-49587, doi:10.1074/jbc.M407340200 (2004).

9 Stevens, J. *et al.* Analysis of the asymmetrically expressed Ablim1 locus reveals existence of a lateral plate Nodal-independent left sided signal and an early, left-right independent role for nodal flow. *BMC Dev Biol* **10**, 54, doi:10.1186/1471-213X-10-54 (2010).

10 Ohsawa, N., Koebis, M., Mitsuhashi, H., Nishino, I. & Ishiura, S. ABLIM1 splicing is abnormal in skeletal muscle of patients with DM1 and regulated by MBNL, CELF and PTBP1. *Genes Cells* **20**, 121-134, doi:10.1111/gtc.12201 (2015).

11 Lahmers, S., Wu, Y., Call, D. R., Labeit, S. & Granzier, H. Developmental control of titin isoform expression and passive stiffness in fetal and neonatal myocardium. *Circ Res* **94**, 505-513, doi:10.1161/01.RES.0000115522.52554.86 (2004).

12 Veerman, C. C. *et al.* Switch From Fetal to Adult SCN5A Isoform in Human Induced Pluripotent Stem Cell-Derived Cardiomyocytes Unmasks the Cellular Phenotype of a Conduction Disease-Causing Mutation. *J Am Heart Assoc* **6**, doi:10.1161/JAHA.116.005135 (2017).

13 Townsend, P. J. *et al.* Human cardiac troponin T: identification of fetal isoforms and assignment of the TNNT2 locus to chromosome 1q. *Genomics* **21**, 311-316, doi:10.1006/geno.1994.1271 (1994).

14 Loiselle, J. J. & Sutherland, L. C. Differential downregulation of Rbm5 and Rbm10 during skeletal and cardiac differentiation. *In Vitro Cell Dev Biol Anim* **50**, 331-339, doi:10.1007/s11626-013-9708-z (2014).

15 Giudice, J. *et al.* Alternative splicing regulates vesicular trafficking genes in cardiomyocytes during postnatal heart development. *Nat Commun* **5**, 3603, doi:10.1038/ncomms4603 (2014).

16 Freiburg, A. *et al.* Series of exon-skipping events in the elastic spring region of titin as the structural basis for myofibrillar elastic diversity. *Circ Res* **86**, 1114-1121, doi:10.1161/01.res.86.11.1114 (2000).

17 Neagoe, C. *et al.* Titin isoform switch in ischemic human heart disease. *Circulation* **106**, 1333-1341, doi:10.1161/01.cir.0000029803.93022.93 (2002).

18 Roof, D. J., Hayes, A., Adamian, M., Chishti, A. H. & Li, T. Molecular characterization of abLIM, a novel actin-binding and double zinc finger protein. *J Cell Biol* **138**, 575-588, doi:10.1083/jcb.138.3.575 (1997).

19 Heinz, S. *et al.* Simple combinations of lineage-determining transcription factors prime cis-regulatory elements required for macrophage and B cell identities. *Mol Cell* **38**, 576-589, doi:10.1016/j.molcel.2010.05.004 (2010).

20 Tabula Muris, C. *et al.* Single-cell transcriptomics of 20 mouse organs creates a Tabula Muris. *Nature* **562**, 367-372, doi:10.1038/s41586-018-0590-4 (2018).

21 D'Antonio-Chronowska, A. *et al.* Association of Human iPSC Gene Signatures and X Chromosome Dosage with Two Distinct Cardiac Differentiation Trajectories. *Stem Cell Reports* **13**, 924-938, doi:10.1016/j.stemcr.2019.09.011 (2019).

22 Tucker, N. R. *et al.* Transcriptional and Cellular Diversity of the Human Heart. *Circulation*, doi:10.1161/CIRCULATIONAHA.119.045401 (2020).

23 Yang, J. *et al.* RBM24 is a major regulator of muscle-specific alternative splicing. *Dev Cell* **31**, 87-99, doi:10.1016/j.devcel.2014.08.025 (2014).

24 Zhang, M. *et al.* Rbm24, a target of p53, is necessary for proper expression of p53 and heart development. *Cell Death Differ* **25**, 1118-1130, doi:10.1038/s41418-017-0029-8 (2018).

25 Tay, J. & Richter, J. D. Germ cell differentiation and synaptonemal complex formation are disrupted in CPEB knockout mice. *Dev Cell* **1**, 201-213, doi:10.1016/s1534-5807(01)00025-9 (2001).

26 Burns, D. M. & Richter, J. D. CPEB regulation of human cellular senescence, energy metabolism, and p53 mRNA translation. *Genes Dev* **22**, 3449-3460, doi:10.1101/gad.1697808 (2008).

27 Bjorkman, K. K., Buvoli, M., Pugach, E. K., Polmear, M. M. & Leinwand, L. A. miR-1/206 downregulates splicing factor Srsf9 to promote C2C12 differentiation. *Skelet Muscle* **9**, 31, doi:10.1186/s13395-019-0211-4 (2019).

28 Stanek, D. *et al.* Prot2HG: a database of protein domains mapped to the human genome. *Database (Oxford)* **2020**, doi:10.1093/database/baz161 (2020).
